# Membrane composition and curvature in SNX9-mediated actin polymerization

**DOI:** 10.1101/2024.09.24.614710

**Authors:** Pankti Vaishnav, Hanae Shimo Kondo, Jonathan R Gadsby, Ulrich Dobramysl, Julia Mason, Joseph Atherton, Jennifer L Gallop

## Abstract

Membrane-binding adaptor protein Sorting nexin 9 (SNX9) contributes to viral uptake and inflammation and is associated with worse outcomes in several cancers. It is involved in endocytosis of epidermal growth factor receptors, β1-integrin and membrane type 1 matrix metalloprotease and in mitochondrial-derived vesicles. Its BAR-PX domain binds phosphatidylinositol phosphates (PIPs) and its SH3 domain interacts with dynamin and N-WASP to stimulate Arp2/3 complex actin polymerization during vesicle scission. Recent complexities have arisen in SNX9’s lipid specificity and its involvement both endocytic and filopodial membrane topologies. Here we use biolayer interferometry, cell-free reconstitution and superresolution microscopy to analyse the activities of SNX9. We find that more SNX9 binds membranes that contain PI(4,5)P2 and PI(3)P compared with PI(3,4)P2, despite having similar affinity, suggesting SNX9 can assemble into different macromolecular arrangements depending on the composition. Actin assembly requires the wider protein and lipid network provided by PX-BAR and SH3 interactions. 3D direct stochastic optical reconstruction microscopy on filopodia-like reconstitutions shows that SNX9 and related protein Transducer of Cdc42 activation-1 (TOCA-1) are competent to form both tubular and plaque-like organizations with the actin machinery. Finally, using cryo-electron tomography we show that SNX9 assembles both branched and bundled actin filaments demonstrating its multifunctional nature.

## Introduction

Sorting Nexin 9 (SNX9) is a multifunctional adaptor protein that plays a critical role in clathrin-mediated endocytosis (Lundmark and Carlsson, 2003), actin dynamics (Yarar *et al*., 2007, 2008; Gallop *et al*., 2013), and membrane remodeling (Pylypenko *et al*., 2007; Shin *et al*., 2008; Hicks *et al*., 2015). SNX9 is also involved in the formation of mitochondria-derived vesicles (MDVs) for mitochondrial quality control and cellular stress responses (Todkar *et al*., 2021; Towers *et al*., 2021; Zecchini *et al*., 2023).

SNX9 expression is increased in metastatic breast cancer cells (Bendris *et al*., 2016b, 2016a), colorectal cancer (Tanigawa *et al*., 2019) and prostate cancer (Shukla *et al*., 2023), amongst others. In metastatic breast cancer, elevated SNX9 levels are associated with enhanced endocytosis of the epidermal growth factor receptor, which drives aggressive cell signaling pathways that promote tumor growth, survival, and invasion. SNX9 expression can also be decreased in primary tumors indicating a context-dependent role (Bendris *et al*., 2016a). In colorectal cancer, SNX9 is upregulated, particularly in advanced stages, correlating with poor prognosis and influencing integrin trafficking, which is essential for cell adhesion, migration, and invasion (Tanigawa *et al*., 2019). In the immune response, SNX9 localizes to the immunological synapse (Ecker *et al*., 2022) and has been implicated in T-cell exhaustion (Trefny *et al*., 2023), inflammation (Ish-Shalom *et al*., 2016) as well as mutations upstream of the SNX9 gene being implicated in increased propensity to developing long Covid (Taylor *et al*., 2023). Because SNX9 deletion in mice has little effect on development or physiology in healthy conditions (Liu *et al*., 2016), SNX9 may represent a disease-relevant target despite its ubiquitous expression across cell types.

Like orthologs SNX18 and SNX33, with which it is partially redundant (Park *et al*., 2010), SNX9 contains conserved PX (phox homology) and BAR (Bin-Amphiphysin-Rvs) domains, which enables it to sense and induce membrane curvature and facilitate vesicle trafficking events. The BAR domain of SNX9 forms a crescent-shaped dimer and the combined BAR-PX domain forms higher order oligomers that can bind a wide variety of phosphoinositide lipids (Pylypenko *et al*., 2007; Yarar *et al*., 2008). The SH3 domain of SNX9 binds Neural Wiskott Aldrich Syndrome Protein (N-WASP), linking the phosphoinositide lipid binding of SNX9 to guanosine triphosphate-bound cell division cycle 42 (Cdc42•GTP) and Actin-related protein (Arp) 2/3 complex-mediated actin polymerization (Yarar *et al*., 2007; Daste *et al*., 2017). SNX9 also interacts with endocytic proteins dynamin and Adaptor Protein-2 (AP-2) (Soulet *et al*., 2005; Lo *et al*., 2017). SNX9 interacts with dynamin, a GTPase essential for vesicle scission, via its proline-rich motifs, allowing SNX9 to coordinate the final steps of vesicle fission through the formation of tight constriction at the vesicle neck (Taylor *et al*., 2012). Binding of clathrin adaptor AP-2 to SNX9 derepresses an autoinhibitory conformation of SNX9, where AP-2 outcompetes the PX-BAR domains in binding to a low-complexity linker region to ensure accurate control of membrane remodelling (Lo *et al*., 2017). Given that around half of clathrin-mediated endocytic events involve N-WASP and actin recruitment in mammalian cells (Taylor *et al*., 2012), actin polymerization linked to endocytosis, scission and subsequent vesicle trafficking represents a key downstream effector function of SNX9.

SNX9 has been reported to bind to a range of phosphoinositide lipids, but which interactions are functionally important is less clear. While the SNX9 PX-BAR domain was reported to bind PI(4,5)P2 and PI(3,4)P2 similarly, the inclusion of additional PI(4,5)P2 binding proteins in computational simulations of endocytosis led to a selectivity of SNX9 for PI(3,4)P2, although the possibility of other PI(3,4)P2 effectors being present was not considered in the models (Schöneberg *et al*., 2017). Although SNX9 is responsible for a 5-fold increase in actin polymerization to membranes containing a coincidence of PI(4,5)P2 and PI(3)P (Gallop *et al*., 2013), it has not been determined how the binding of SNX9 to PI(4,5)P2/PI(3)P compares to PI(3,4)P2.

SNX18 and SNX33 also bind well to PI(4,5)P2/PI(3)P and mutations in the BAR domain reduces SNX9 binding to PI(4,5)P2 and mutations in the PX domain to PI(3)P respectively (Daste *et al*., 2017). Consistent with a functional role for PI(3)P, reducing the levels of inositol polyphosphate 4-phosphatase (INPP4A) that hydrolyses PI(3,4)P2 to PI(3)P reduced endocytosis in HeLa cells (Daste *et al*., 2017) and an auxilin conjugated-PI(3)P probe localizes to endocytic sites (He *et al*., 2017). Either PI(3)P or PI(3,4)P2 can alleviate the inhibition in endocytosis caused by reducing the levels of class II phosphoinositide 3 kinase (PI3K), again supporting a functional role for PI(3)P, though a role during endocytosis for the relevant 4-phosphatases INPP4A/B that could produce PI(3)P from PI(3,4)P2 is not as clear as its later role at endosomes (Wang *et al*., 2018).

The picture of how SNX9 operates in cells is complicated by the involvement of SNX9 in negative membrane curvature scenarios. The BAR-PX domain bends membranes in positive (endocytic) directions yet *Chlamydia trachomatis* uses SNX9 to increase viral spread by filopodia (Ford *et al*., 2018), finger-like protrusions from the cell surface comprising a bundle of actin filaments surrounded by highly negatively curved plasma membrane. This suggests that SNX9 can play important roles at sites of actin polymerization regardless of the apparent curvature preference of the BAR-PX domain, perhaps due to actin forces overriding this preference. Consistent with this, overexpression of SNX9 causes filopodial protrusion in mammalian and *Drosophila* cells (Hicks *et al*., 2015; Bendris and Schmid, 2017) and endogenous SNX9 localises to filopodial tips in cultured cells and a subset of *Xenopus* cell filopodia (Jarsch *et al*., 2020). Other positively curved BAR domain proteins, including transducer of Cdc42-dependent actin assembly protein 1 (TOCA-1), localise to filopodia (Saengsawang *et al*., 2012; Taylor *et al*., 2019; Blake *et al*., 2024) and it is not known how the BAR domain curvature is reconciled with the local membrane curvature at these sites. SNX9 is linked to the formation of branched actin via N-WASP and the Arp2/3 complex, while TOCA-1, also interacts with Drf3, Ena and VASP to promote linear actin bundles. It is not clear whether SNX9 can contribute to making bundled actin, as implied by its filopodia localization.

We recently found that our cell-free system of filopodia-like structures (FLS) requires SNX9, and that metabolism of PI(4,5)P2 through PI3K activity to PI(3)P was essential for its involvement (Ford *et al*., 2018; Jarsch *et al*., 2020). This system uses *Xenopus* egg extracts, subjected to a high speed centrifugation step, which give a complex cytosolic environment without endogenous membranes. When applied to supported lipid bilayers enriched with 10% mole fraction PI(4,5)P2, these extracts reconstitute filopodia-like structures that grow up from the glass coverslip. They are visualized using fluorescently labelled G-actin, or other fluorescent probes added into the extracts, and spinning disk confocal microscopy (Lee *et al*., 2010). Here, we sought to shed light on how SNX9 functions in complex environments by comparing the response of purified SNX9 to PI(4,5)P2/PI(3)P and PI(3,4)P2, and by examining the role of SNX9 domains, interaction partners and topologies of the actin and membrane structures associated with SNX9 in the cell-free system.

## Results

### The binding characteristics of SNX9 to membranes of different composition implies different multimeric arrangements

SNX9 is frequently described as a PI(3,4)P2 binding protein (Posor *et al*., 2013; Schöneberg *et al*., 2017). Similar binding of purified SNX9 to PI(3,4)P2 as to PI(4,5)P2 has been seen by surface plasmon resonance (Schöneberg *et al*., 2017), similar to earlier sedimentation assays that showed promiscuous SNX9 binding to different phosphoinositides (Pylypenko *et al*., 2007; Yarar *et al*., 2008). However, for its activities in actin polymerization *in vitro*, PI(4,5)P2 with PI(3)P is the far superior lipid combination (Gallop *et al*., 2013; Daste *et al*., 2017). Phosphoinositide lipids can be interconverted, which complicates data interpretation (Figure 1A).

**Figure 1.**
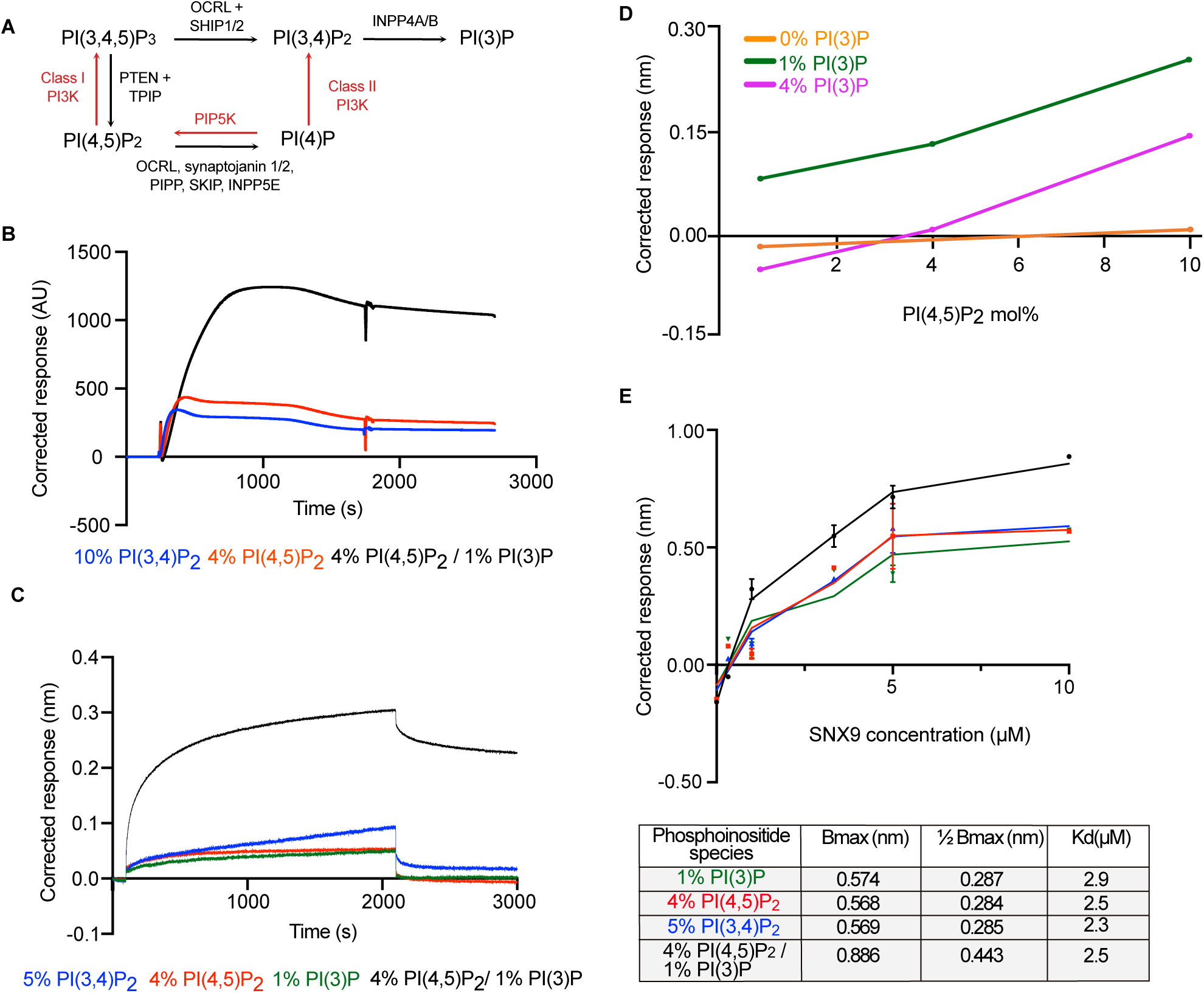
Greater extent while similar half maximal SNX9 binding to PI(4,5)P2 / PI(3)P compared to PI(3,4)P2 or P(4,5)P2. (A) Schematic of phosphoinositide lipid conversion (B) SPR trace of at 1 µM protein SNX9 binding to liposomes containing 60-70% PC, 30% PS and the indicated phosphoinositide lipid shows more binding to PI(4,5)P2/PI(3)P. (C) BLI trace of SNX9 binding to liposomes containing 65-70% PC, 30% PS and the indicated phosphoinositide lipid at 1 µM protein. (D) There is an optimum of SNX9 binding at 1% mol fraction PI(3)P with dose-dependent binding to PI(4,5)P2. (E) Summary response of all phosphoinositide lipids to SNX9 at increasing protein concentrations.

To address these contradictions, we compared the binding of SNX9 to membranes of different composition using surface plasmon resonance (SPR) and biolayer interferometry (BLI). We used a membrane composition similar to the inner leaflet of the plasma membrane. To maximise membrane integrity we used 60-70% phosphatidylcholine (PC, rather than phosphatidylethanolamine) and 30% phosphatidylserine to retain a representative level of plasma membrane negative charge. We then substituted PC for the phosphoinositide lipids. Liposomes comprised of 70% PC/30% PS were included in each experiment and the trace subtracted from the phosphoinositide lipid-containing liposome trace. Occasionally the PC/PS trace was aberrantly high, and in that case if the phosphoinositide runs had been consistent with previous observations, we used the average of 9 other control runs for background subtraction.

Using 1 µM full-length SNX9, the presence of both 4% PI(4,5)P2 and 1% PI(3)P together in the liposomes gives substantially increased SNX9 binding compared with PI(3,4)P2 or PI(4,5)P2 alone (Figure 1B, by SPR, recapitulated with BLI Figure 1C). Binding increases over the first 600 s and reaches a near-plateau by 2000 s. Little dissociation is observed in any condition. We found that BLI was less complicated to implement as in SPR the injection needle frequently clogged in the absence of detergents. Therefore, we continued with BLI. To assess the relative contribution of PI(4,5)P2 and PI(3)P, we titrated the mole fraction of either PI(4,5)P2 or PI(3)P whilst keeping the other constant. Existing binding models were a poor fit to the data so we quantified the overall response by taking the average response in the last 20 s of the association phase (Figure 1D, with the original background-subtracted traces in Sup. Figure 1A-C). In the absence of PI(3)P, the binding to PI(4,5)P2 alone is no greater than binding to the PC/PS background (Figure 1D, orange line, Sup Figure 1A). Addition of 1% PI(3)P leads to increases in binding across 1%, 4% and 10% PI(4,5)P2 (Figure 1D, green and magenta lines, Sup. Figure 1B). Unexpectedly, a further increase to 4% PI(3)P decreases binding, suggesting that there is an optimum density of PI(3)P for the SNX9 interaction (Figure 1D magenta line compared with green line, Sup. Figure 1C). As BAR domain proteins form multimeric protein arrangements on the membrane surfaces, this suggests that the SNX9 protein-protein and protein-lipid interactions involved can be disrupted by different balances of phosphoinositide lipid interactions.

We then measured binding at progressive concentrations of SNX9 at the different key membrane compositions (summary data Figure 1E, original traces Sup. Figure 1D-G). Taking PI(4,5)P2/1% PI(3)P as our reference, we measured the response to increasing SNX9 at 4% PI(4,5)P2, 1% PI(3)P and 5% PI(3,4)P2. Overall, PI(4,5)P2 / PI(3)P gives more binding at all SNX9 concentrations (Figure 1E).

The behaviours are also complex as the shapes of the original traces show differences between lipid compositions and the nature of the binding changes with protein concentration (Sup. Figure 1E-G). The binding curve to PI(4,5)P2 and PI(3)P alone first rises quickly and then continues a slow rise and does not complete binding over 2000 s (Sup. Figure 1D,F). In contrast, PI(3,4)P2 and PI(4,5)P2/PI(3)P rise quickly and have a smooth transition to a phase of gradually increasing binding (Sup Figure 1E, G). While 1 µM SNX9 gives a plateau when interacting with 4% PI(4,5)P2 / 1% PI(3)P (Sup. Figure 1G) at higher SNX9 concentrations binding continues to rise, showing a trend towards a plateau with PI(3,4)P2 at 10 µM SNX9 and not with the others (Sup. Figure 1E). In all cases, there is little dissociation, suggesting a strong matrix of protein-protein and protein-lipid interactions.

The response of SNX9 binding to protein concentration to PI(3,4)P2 and PI(4,5)P2 is quantitatively similar to previous measurements using the BAR-PX domain (Lo *et al*., 2017; Schöneberg *et al*., 2017; Chandra *et al*., 2019). The slightly higher half maximal binding we observe (2.5 µM SNX9 compared to 1.5 µM in previous work) could arise from the autoinhibitory interaction between the SH3 domain and the BAR domain because we used full-length protein. The data indicate that SNX9 binds to PI(4,5)P2 plus PI(3)P with the same affinity as PI(4,5)P2 or PI(3,4)P2 alone, but interestingly, with PI(4,5)P2 plus PI(3)P the extent of SNX9 binding is much greater (Figure 1E). Together with the distinct shapes of the binding curves (Sup Figure 1D-G), we think this indicates that there is a different multimeric arrangement of SNX9 taking place on membranes of different composition.

### Within a cytosolic environment PI(3)P increases actin assembly and SNX9 recruitment to actin sites whereas PI(3,4)P2 does not

To visualize the contributions of PI(3)P concentration to SNX9 activity at actin superstructures, we used the cell-free system of FLS that nucleate on PI(4,5)P2-enriched supported lipid bilayers (Lee *et al*., 2010). We visualized actin using Alexa488-conjugated rabbit actin and labelled SNAP-SNX9 with chemical fluorophore Alexa Fluor-647 *in vitro*, titrating it into the extracts at low levels that did not affect FLS growth. FLS have the advantage that while the actin is dynamic, the membrane clustered complex of proteins is anchored near the glass and the proteins that cluster to the actin sites can be quantified.

We examined the dose-dependent response to PI(3)P by adding it in the presence and absence of wortmannin, which will inhibit phosphorylation of PI(4,5)P2 by the extracts (Figure 2). As described previously, wortmannin treatment eliminates FLS (Jarsch *et al*., 2020). We saw a dose-responsive increase in FLS with increasing mole fraction of PI(3)P in the presence of wortmannin, suggesting that the presence of PI(3)P is the limiting component. Without wortmannin no further activation or inhibition is observed suggesting that native levels of phosphoinositide conversion in the extracts are making the PI(3)P already (Figure 2A-C). These results also suggest that the inhibition by PI(3)P observed at high levels by BLI may not be relevant in a more complex protein milieu where SNX9 is binding other proteins and other proteins will also be binding the PI(3)P. Higher levels of PI(3)P increased SNX9 intensity at FLS while the fraction of FLS with SNX9 present above background intensity levels stayed at approximately 30% (Figure 2D-E).

**Figure 2.**
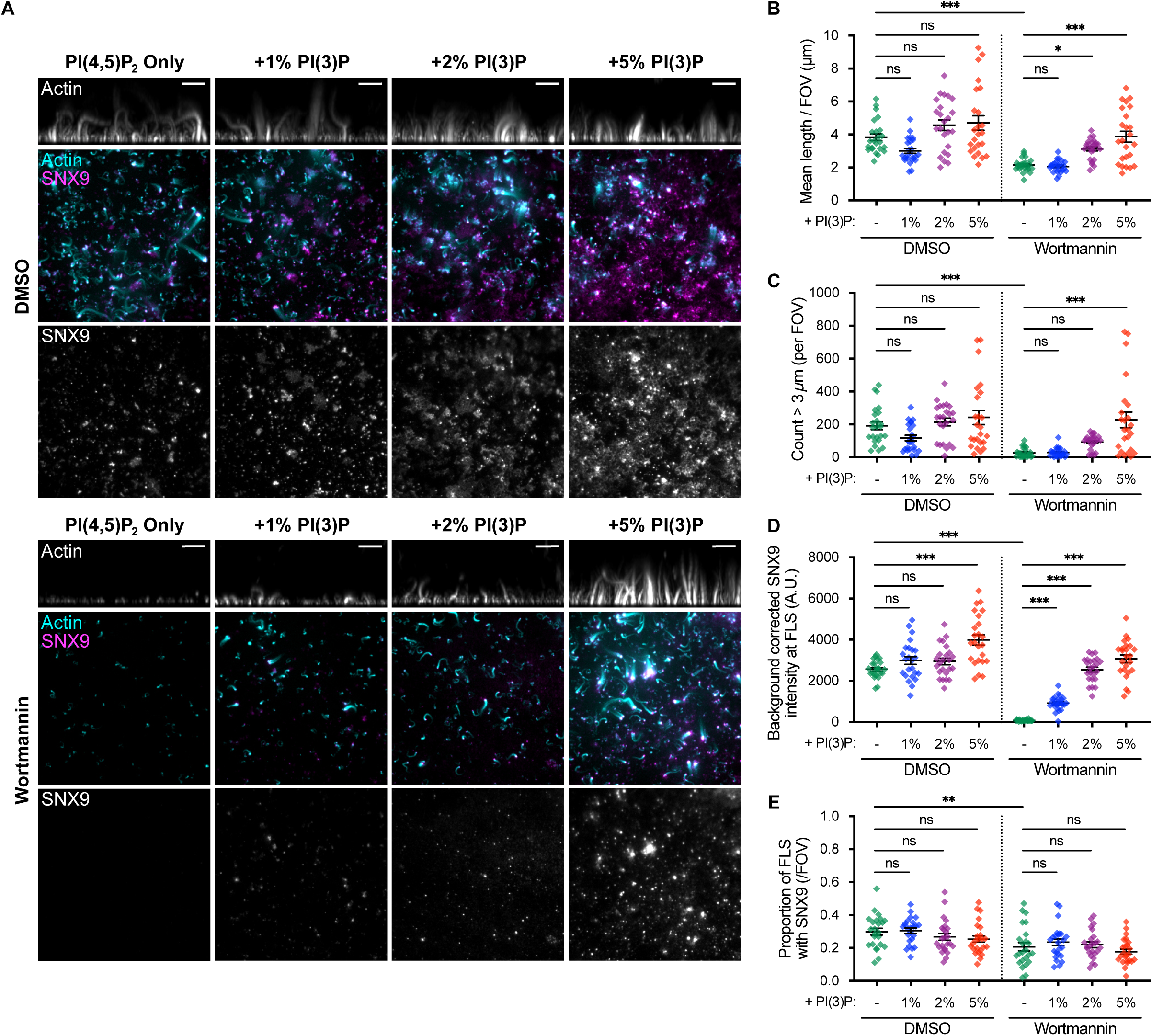
Increased FLS length and numbers driven by PI(3)P is associated with increased levels of SNX9 localizing to FLS. (A) TIRF and confocal imaging of representative fields of view (FOV) of FLS grown from membranes supplemented with PI(3)P as indicated, in the presence or either 2 µM wortmannin or the DMSO vehicle control. The upper image is a maximally projected side view of a 30 x 1 micron confocal stack of the actin channel, while the middle composite image shows a maximal Z projection of the actin confocal stack (cyan) and a TIRF image of SNX9 at the coverslip surface (magenta, and lower single channel image). Scale bars = 10 µm. (B-E) Quantification of (B) mean actin structure length per field of view, (C) count of FLS > 3 µm, (D) SNX9 intensity at FLS and (E) the proportion of LFS containing SNX9. In each quantification, each datapoint represents an individual FOV, lines indicate mean ± SEM, N = 24 FOVs. Statistical significance for each was assessed by ordinary one-way ANOVA with Holm-Sidak’s multiple comparisons test. Overall ANOVA for each quantification p < 0.001. Individual comparisons: (B) DMSO PI(4,5)P2 only vs DMSO 1% p = 0.06, vs DMSO 3% p = 0.07 & vs DMSO 5% p = 0.05 . DMSO PI(4,5)P2 only vs wortmannin PI(4,5)P2 only = p < 0.001. Wortmannin PI(4,5)P2 only vs wortmannin 1% p = 0.84, vs wortmannin 3% p = 0.02, vs wortmannin 5% p < 0.001. (C) DMSO PI(4,5)P2 only vs DMSO 1% p = 0.23, vs DMSO 3% p = 0.79 & vs DMSO 5% p = 0.45. DMSO PI(4,5)P2 only vs wortmannin PI(4,5)P2 only p < 0.001. Wortmannin PI(4,5)P2 only vs wortmannin 1% p = 0.95, vs wortmannin 3% p = 0.30 [both n.s.], vs wortmannin 5% p < 0.001. (D) DMSO PI(4,5)P2 only vs DMSO 1% p = 0.10, vs DMSO 3% p = 0.10 [both n.s], vs DMSO 5% p < 0.001. DMSO PI(4,5)P2 only vs Wortmannin PI(4,5)P2 only p < 0.001. Wortmannin PI(4,5)P2 only vs wortmannin 1%, vs wortmannin 3% & vs wortmannin 5% all p < 0.001. (E) DMSO PI(4,5)P2 only vs DMSO 1% p = 0.87, vs DMSO 3% p = 0.80 & vs DMSO 5% p = 0.50. DMSO PI(4,5)P2 only vs wortmannin PI(4,5)P2 only p = 0.009. wortmannin PI(4,5)P2 only vs wortmannin 1% p = 0.80, vs wortmannin 3% p = 0.87 & vs wortmannin 5% p = 0.80.

We have previously observed that PI(3,4)P2 does not functionally rescue in the presence of wortmannin (Daste *et al*., 2017). Therefore, we compared the supplementation of PI(3,4)P2, by comparison to PI(3)P in native conditions (Figure 3A-B). There is a significant decrease in FLS length with additional PI(3,4)P2 compared with PI(4,5)P2 alone or supplemented PI(3)P (Figure 3C). In liposomes adding PI(3,4)P2 let to a decrease in polymerization when no PI(3)P was present by pyrene actin measurements as well (Daste *et al*., 2017). Both supplemented conditions slightly decrease the numbers of FLS forming on the membrane, perhaps due to other binding proteins being recruited to the membrane, occluding possible sites of FLS formation (Figure 3D).

**Figure 3.**
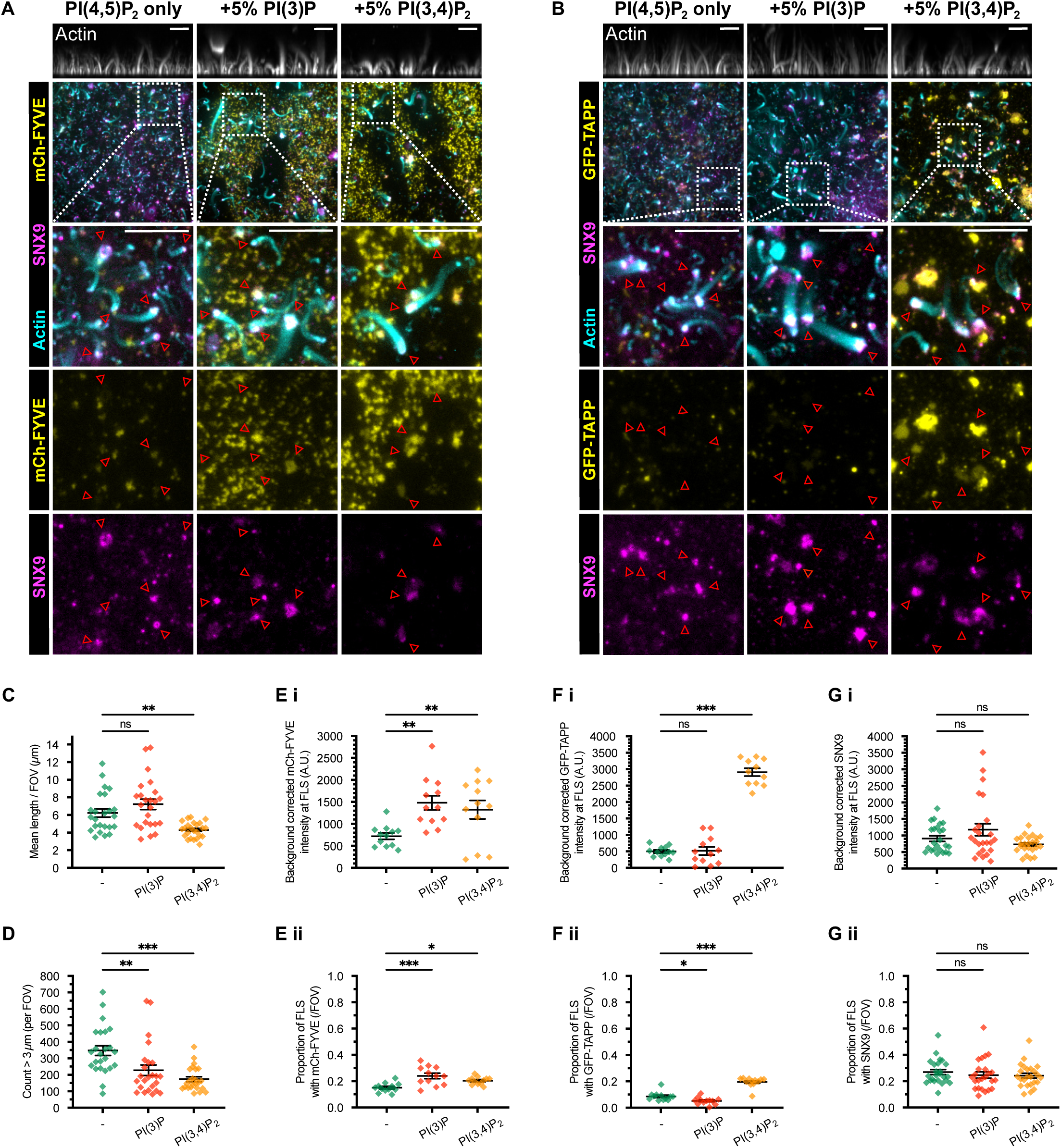
There are fewer and shorter FLS when PI(3,4)P2 is added to supported lipid bilayers, which causes both increased PI(3,4)P2 and PI(3)P probe localization. (A-B) TIRF and confocal imaging of representative fields of view (FOV) of FLS grown from membranes supplemented with 5% PI(3)P or PI(3,4)P2, 2xFYVE to detect PI(3)P and 2xPH(TAPP)(Posor *et al*., 2013) to detect PI(3,4)P2 as indicated. The upper image is a maximally projected side view of a 30 µm confocal z-stack in 1 µm increments, of the actin channel, while the image shows composite of a maximal z projection in the x-y plane of the actin confocal stack (cyan) and TIRF images at the coverslip surface of SNX9 (magenta) and either (A) mCh-FYVE or (B) GFP-TAPP (yellow). The boxed region is enlarged in the lower three images illustrated the three colour composite and SNX9, mCh-FYVE or GFP-TAPP channels separately, as indicated. All scale bars = 10 µm. On the enlargements several FLS are marked (red arrows) to allow comparison of SNX9, mCh-FYVE or GFP-TAPP localization under FLS, (C-G) Quantification of (C) mean FLS length per field of view, (D) count of FLS > 3 µm, (E) mCherry-FYVE (i) intensity and (ii) proportion of labelled FLS, (F) GFP-TAPP (i) intensity and (ii) proportion of labelled FLS, and (G) SNX9 (i) intensity and (ii) proportion of labelled FLS, In each quantification, each datapoint represents an individual FOV, lines indicate mean ± SEM, For (C, D & G), N = 24 FOVs, For (E & F) N = 12 FOVs. Statistical significance for each was assessed by ordinary one-way ANOVA with Holm-Sidak’s multiple comparisons test. Overall ANOVA for (D, D, Ei, Fi, Fii & Gi) p < 0.001, for [Eii] p = 0.003, for [Gi] p = 0.006, for [Gii] p = 0.12 [ns]. Individual comparisons: (C) PI(4,5)P2 only vs + PI(3)P p = 0.19, vs + PI(3,4)P2 p = 0.005. (D) PI(4,5)P2 only vs + PI(3)P p = 0.006, vs + PI(3,4)P2 p < 0.001. (Ei) PI(4,5)P2 only vs + PI(3)P p = 0.002, vs + PI(3,4)P2 p = 0.007. (Eii) PI(4,5)P2 only vs + PI(3)P p < 0.001, vs + PI(3,4)P2 p = 0.02. (Fi) PI(4,5)P2 only vs + PI(3)P p = 0.92, vs + PI(3,4)P2 p < 0.001. (Fii) PI(4,5)P2 only vs + PI(3)P p = 0.03, vs + PI(3,4)P2 p < 0.001. (Gi) PI(4,5)P2 only vs + PI(3)P p = 0.22, vs + PI(3,4)P2 p = 0.25. (Gii) PI(4,5)P2 only vs + PI(3)P p = 0.52, vs + PI(3,4)P2 p = 0.52.

To visualize the locations of added phosphoinositide lipids in this experiment we employed PI(3)P binding probe mCherry-2xFYVE (Gillooly *et al*., 2000) and PI(3,4)P2 binding probe eGFP-PH2x(TAPP) (Dowler *et al*., 2000; Posor *et al*., 2013). The intensity of 2xFYVE detected in the supported bilayers under FLS is increased with both PI(3)P and PI(3,4)P2, as well as the proportion of FLS with 2xFYVE localization, showing that PI(3)P can be produced at FLS from the added PI(3,4)P2 (Figure 3E). The intensity of 2xTAPP is low in native conditions and increases in both intensity and proportion of FLS when additional PI(3,4)P2 is added and a small decrease when PI(3)P is added (Figure 3F). Thus, PI(3,4)P2 is not being made from PI(3)P. The SNX9 intensity and proportion of labelled FLS remained similar across the supplemented conditions again confirming that native levels provided by the extract enzyme activities are sufficient (Figure 3G).

Overall, our data indicates that in the presence of PI(4,5)P2, PI(3)P stimulates both SNX9 recruitment and resulting actin polymerization whereas PI(3,4)P2 is inhibitory, despite its similar binding affinity to SNX9. Combined with the BLI data, we suggest that the multimeric arrangement attained by SNX9 on PI(4,5)P2 /PI(3)P membranes is competent for actin polymerization and on PI(3,4)P2 membranes it is not.

### In a complex cytosolic environment, the different SNX9 domains and major interaction partners are all needed for actin assembly

Having looked at the lipid compositions, we then looked at the contributions of the different domains of SNX9 to FLS growth. To do this, we immunodepleted SNX9 from the extracts (Figure 4A) and added back purified full-length SNX9 with point mutations in the BAR domain (K511E/K517E), PX domain (Y276A/K302A) or SH3 domain (W42K) (the protein purification gels are shown in Sup. Figure 2). The BAR and PX domain mutations both reduce SNX9 binding to PI(4,5)P2 + PI(3)P by BLI (Sup. Figure 2). All three mutated proteins are unable to restore either FLS number, length or base area compared to re-addition of wild type SNX9 (Figure 4 C-E). This demonstrates that all three interactions of the BAR domain with PI(4,5)P2, the PX domain with PI(3)P and the SH3 domain with N-WASP and other binding partners are essential for its activity (Figure 4B-E).

**Figure 4.**
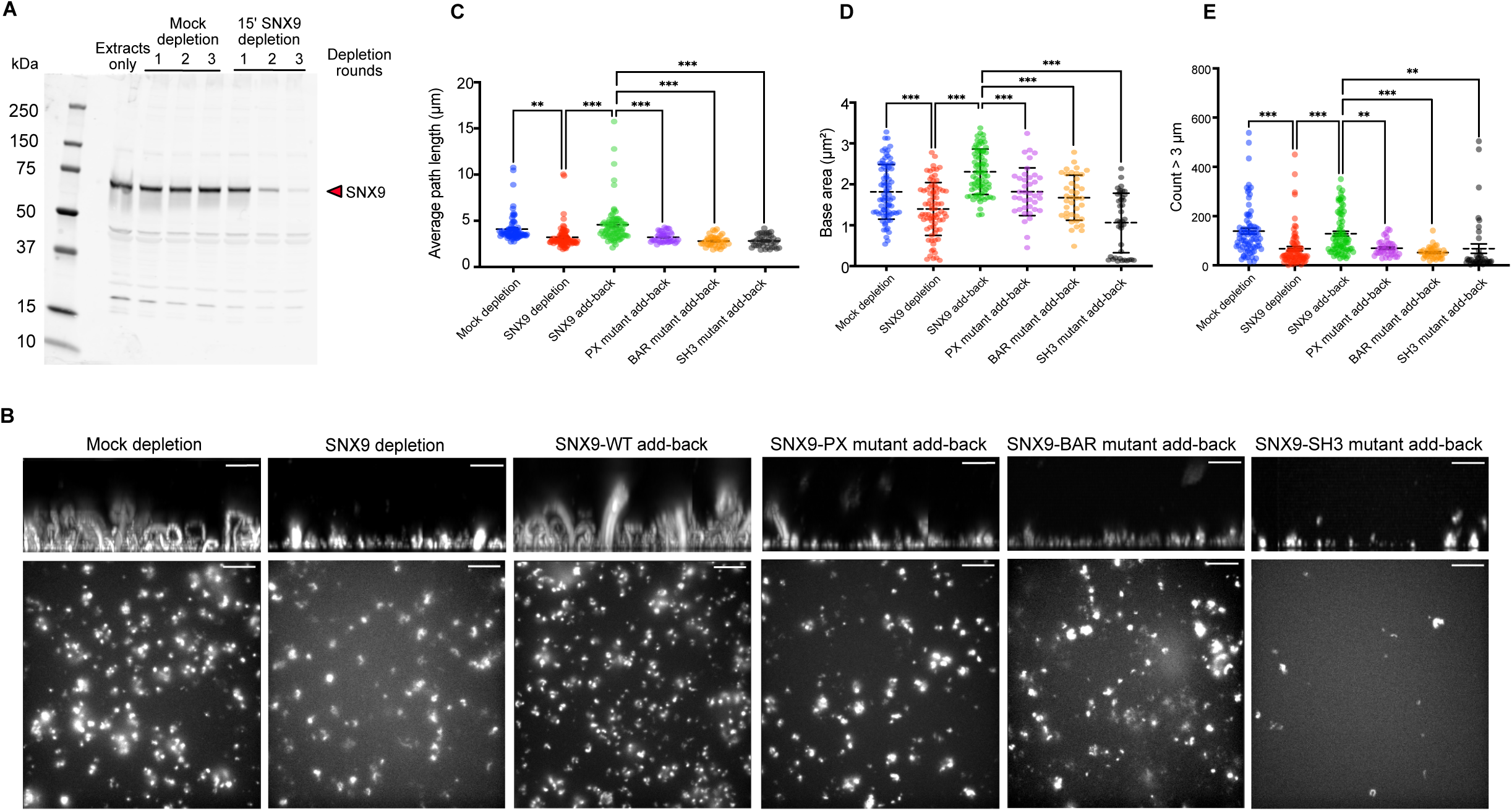
All SNX9 domains are required for actin assembly in FLS. (A) Western blot staining SNX9 showing immunodepletion of SNX9 from HSS. Faint bands are at the molecular weights of antibody heavy and light chain. (B) Confocal imaging of representative fields of view (FOV) of FLS grown from either mock depleted or SNX9 depleted alone or with added indicated SNX9 mutants. The residue mutations were BAR domain: K511E/K517E, PX domain: Y276A/K302A and SH3 domain: W42K (C) Numbers of FLS growing beyond 3 µm are reduced with SNX9 depletion and restored with SNX9 and not with any of the mutants. Statistical significance for each was assessed by ordinary one-way ANOVA with Holm-Sidak’s multiple comparisons test. Overall ANOVA P values: mock vs SNX9 depletion: p <0.001, SNX9 depletion vs SNX9 add back: p <0.001, SNX9 vs PX mutant add back p = 0.004, SNX9 vs BAR mutant add back: p <0.001, SNX9 vs SH3 mutant add back: p = 0.003. (D) Similar to (C) measuring length. Mock vs SNX9 depletion: p = 0.001, SNX9 depletion vs SNX9 add back: p <0.001, SNX9 vs PX add back: p <0.001, SNX9 vs BAR add back: p <0.001, SNX9 vs SH3 add back: p <0.001(E) Base area of the FLS measured with actin gives similar results. Mock vs SNX9 depletion: p <0.001, SNX9 depletion vs SNX9 add back: p <0.001, SNX9 vs PX add back: p <0.001, SNX9 vs BAR add back: p <0.001, SNX9 vs SH3 add back: p <0.001.

It is known that immunodepletion of N-WASP inhibits FLS (Lee *et al*., 2010). As well as N-WASP, the SH3 domain of SNX9 notably interacts with dynamin which is implicated in bundling actin independently of its role in endocytosis (Zhang *et al*., 2020). We found that dynamin localizes to FLS similar to previous data with *Listeria* actin comets (Figure 5A-C, (Lee and Camilli, 2002)), whereas endocytic proteins endophilin and clathrin do not co-localize with actin (Figure 5B). This is likely because there are no transmembrane proteins within the supported lipid bilayers. To test the functional role of dynamin we used dynole compounds, where 34-2 is active and 31-2 an inactive chemically related compound (Hill *et al*., 2009). Dynole 34-2 inhibits FLS formation and growth (Figure 5C-E) and also prevents SNX9 and dynamin recruitment to the membrane (Figure 5F-G). These data again demonstrate that the interaction of SNX9 with multiple binding partners is critical for actin assembly at FLS.

**Figure 5.**
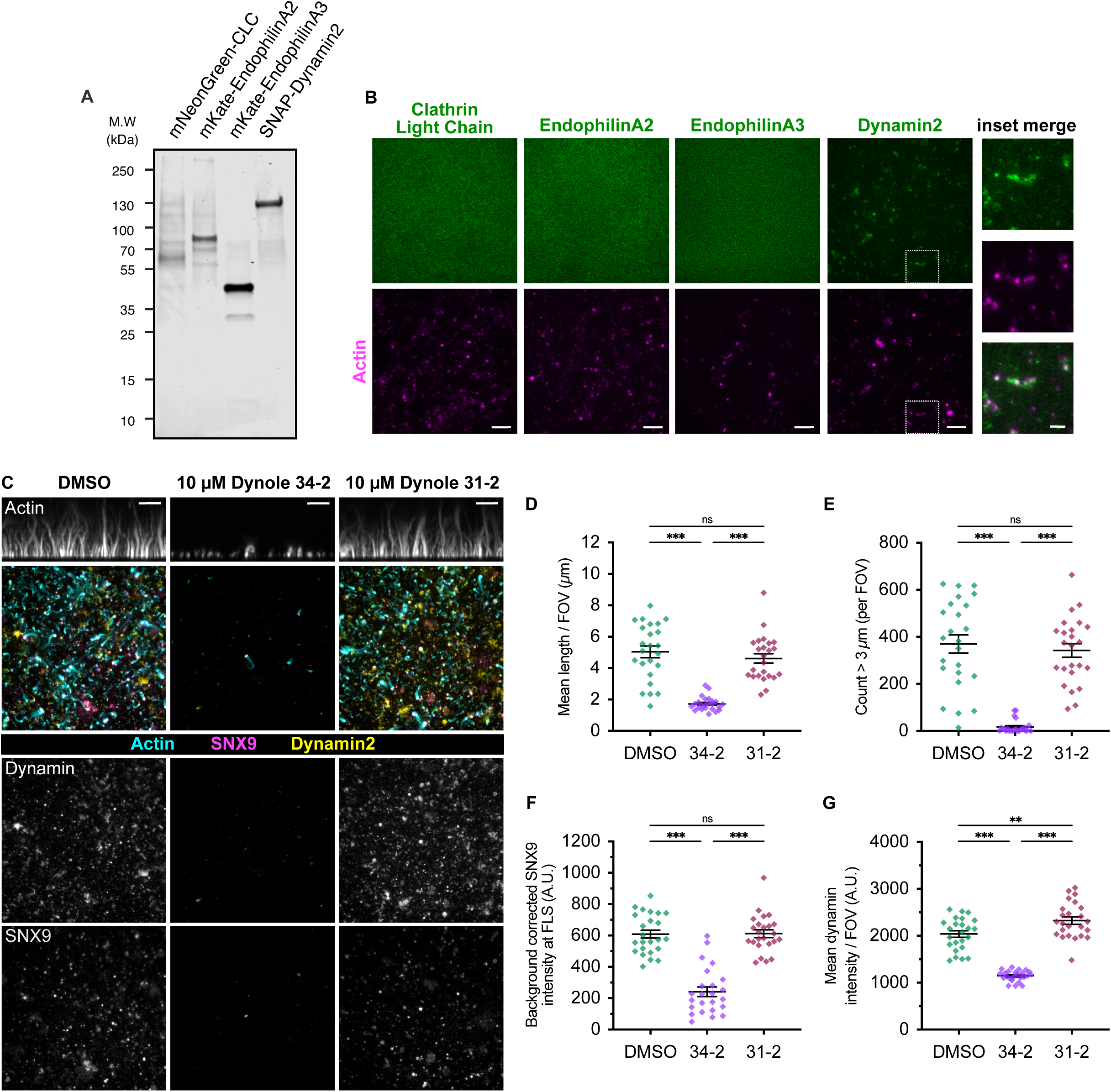
Dynamin 2 localizes to FLS unlike endocytic proteins clathrin and endophilin and FLS are inhibited by active Dynole dynamin inhibitor. (A) Coomassie stained SDS-PAGE gel of purified mNeon-Green Clathrin heavy chain, mKate-EndophilinA2, mKate-Endophilin A3 and SNAP-Dynamin2. (B) TIRF imaging of representative fields of view (FOV) of FLS grown with purified mNeon-Green Clathrin heavy chain, mKate-EndophilinA2, mKate-Endophilin A3 and Alexa-488 labelled SNAP-Dynamin2, as indicated showing dynamin localization only. Actin growth in z imaged by spinning disk confocal. Scale bars = 10 µm. Insets show dynamin 2 and actin merge, scale bar = 3 µm (C) TIRF and confocal imaging of representative fields of view (FOV) of FLS grown in the presence or either 10 µM Dynole 34-3, 31-2 or the DMSO vehicle control, as indicated. The upper image is a maximally projected side view of a 30 x 1 micron confocal stack of the actin channel, while the composite image below shows a maximal Z projection of the actin confocal stack (cyan) and TIRF images of SNX9 (magenta, upper single channel image) and Dynamin 2 (yellow, lower single channel image) at the coverslip surface. Scale bars = 10 µm. (D-G) Quantification of (D) mean actin structure length per field of view, (E) count of FLS > 3 µm, (F) SNX9 intensity at FLS and (G) mean dynamin-2 intensity across a FOV. In each quantification, each datapoint represents an individual FOV, lines indicate mean ± SEM, N = 24 FOVs. Statistical significance for each was assessed by ordinary one-way ANOVA with Holm-Sidak’s multiple comparisons test. Overall ANOVA for each quantification p < 0.001. Individual comparisons: (D) DMSO vs 31-2 & 34-2 vs 31-2, both p <0.001, DMSO vs 31-2 p = 0.29 [ns]. (E) DMSO vs 31-2 & 34-2 vs 31-2, both p <0.001, DMSO vs 31-2 p = 0.49, (F) DMSO vs 31-2 & 34-2 vs 31-2, both p <0.001, DMSO vs 31-2 p = 0.93. (G) DMSO vs 31-2 & 34-2 vs 31-2, both p <0.001, DMSO vs 31-2 p = 0.002.

### In a complex cytosolic environment, SNX9 and related F-BAR domain protein TOCA-1 can form distinct architectures in collaboration with actin and other components

The final aspect of SNX9 interactions we considered is the membrane topology. SNX9 forms positively-curved tubules of the plasma membrane in cells as well as *in vitro* using its BAR domain, although it can also form filopodial protrusions (Bendris and Schmid, 2017). To visualize the topology of SNX9 clusters at FLS we used 3D stochastic optical reconstruction microscopy (STORM) using a total internal reflection fluorescence (TIRF) microscope fitted with a cylindrical lens to assign positions in the z-axis. FLS were visualized in parallel with Alexa Fluor-488 actin imaged by wide field illumination. We subjected the field of view to repeated switching of the Alexa 647 fluorophore labelling either SNX9 or TOCA-1 and reconstructed STORM images.

Typically 6-7 SNX9 foci underneath actin-rich FLS could be seen in any one field of view (Figure 6A). While by TIRF microscopy the foci are indistinguishable (Figure 6B), by STORM three broad types of SNX9 arrangement could be seen: flat, plaque-like clusters that do not elevate in the z-axis (a, b), tubule-like with a lumen empty of SNX9 localizations (c, d, e) and a hybrid of both (f, g). An elevation of 500 nm is typically observed, which was within the calibration range (Figure 6C). The x-y width of the tubules is around 1 µm with the plaques typically wider, so overall a similar membrane area may be covered (Figure 6D). To test whether a related protein at FLS shows similar arrangements, we examined F-BAR domain protein Transducer of Cdc42 activation 1 (TOCA-1) and saw equivalent results (Figure 6E-H).

**Figure 6.**
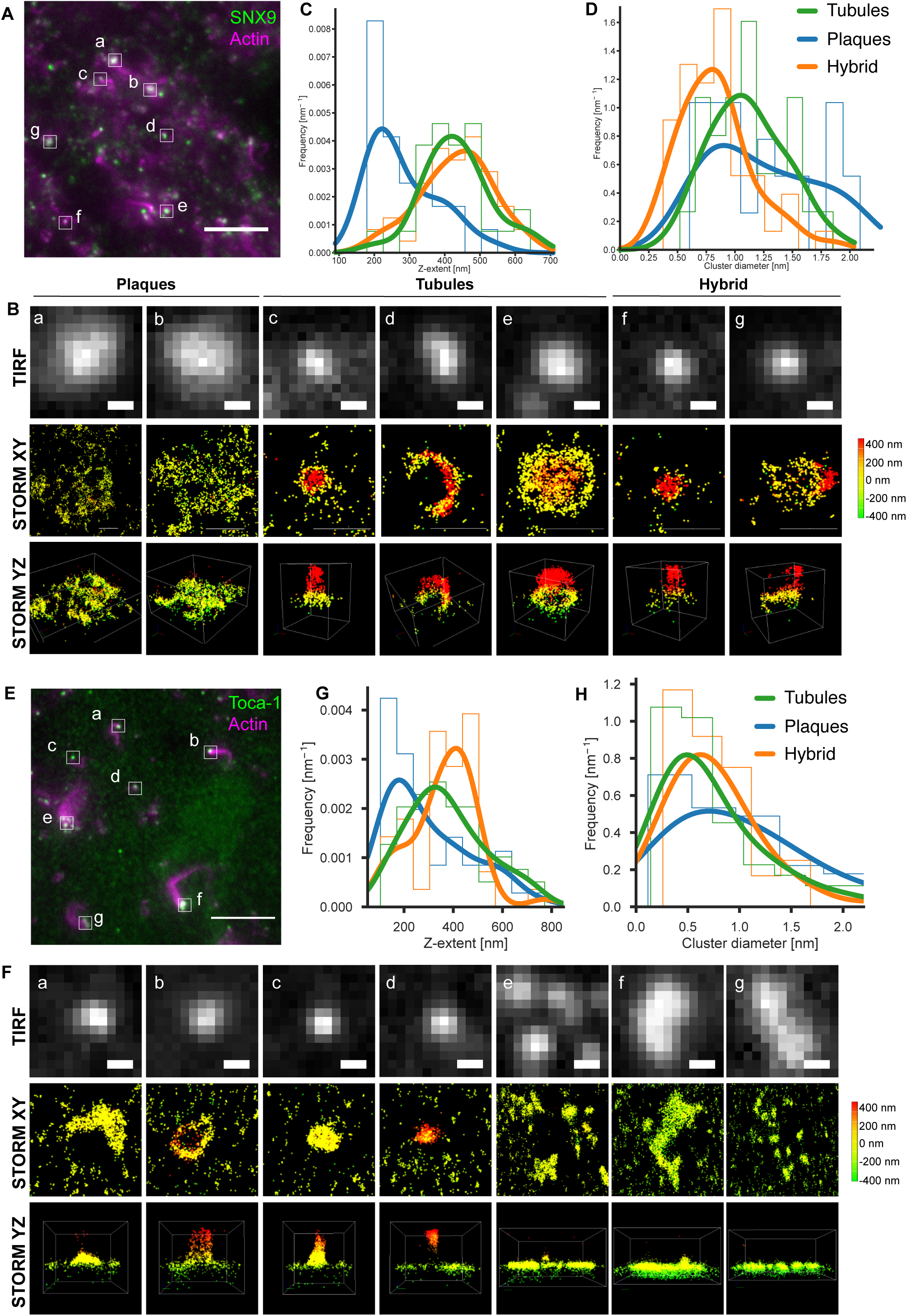
3D STORM shows tubular, plaque-like and hybrid morphologies of SNX9 at FLS sites. (A) Representative microscopy image showing SNX9 (green) by TIRF and actin by widefield (magenta). Scale bar 10 µm. (B) Closeup and STORM images of indicated FLS showing plaque-like (a, b), tubular (c, d, e) and hybrid morphologies (f, g). The first row is the TIRF image with no discernable structural features. The second and third rows represent lateral and axial view of the 3D STORM reconstructions. Depth in the z-plane is colour coded from red (+400 nm) to green (-400 nm). Scale bar 0.5 µm. (C) Numbers of each morphology with given z-extent displayed as a frequency diagram. (D) Similar plot for the diameter of the localization areas.

We wondered whether the tubule structures could be caused by liposomes remaining on the supported bilayer surface, leading to nucleation of the FLS. To test this we labelled PLCδ PH domain, applied it to the supported bilayers without frog egg extract and performed a STORM experiment. No vesicles or elevations similar to those seen with SNX9 could be observed (Sup. Figure 3A). Adding both PLCδ PH domain and TOCA-1 again failed to lead to tubule or plaque formation, which were only formed when extracts were added (Sup. Fig 3B,C). Tubules and plaques could also be seen without the fixation step (Sup. Figure 3D,E). The presence of tubules and plaques was independent of the TOCA-1 concentration added in (Sup. Figure 4). To test the contributions of the extracts, we used different concentrations. At higher extract concentrations, FLS are shorter and wider, presumably due to differences in the balance between disassembly and assembly factors (Sup. Figure 5A-C). More tubules are seen at lower extract concentrations, when the FLS growing from them is longer (Sup. Figure 5F-G).

To test whether the tubule and plaque structures were formed by actin polymerization, we added actin monomer-sequestering drug latrunculin B (LatB) to the extracts and performed STORM experiments on Alexa-647 SNX9 and TOCA-1, as before. While tubules and plaques are both still evident (Figure 7B), the surface area and height covered by the localizations is reduced (Figure 7C-D). The number of localizations of SNX9 has a strong reduction in the presence of LatB (Figure 7E). The TOCA-1 foci present form both tubule and plaque morphologies and there is less effect of LatB on the z-extent or cluster diameter (Figure 7F-G). There is also less reduction in TOCA-1 localizations consistent with previous observations showing TOCA-1 remains on the membrane with LatB treatment (Dobramysl *et al*., 2021) (Figure 7H-J). The data indicates that BAR domain proteins collaborate with multiple other proteins present in the extracts to assemble into either tubules or plaques on the PI(4,5)P2 supported bilayers. In addition, the actin-driven network is important for SNX9 recruitment (consistent with the SH3 domain mutant result in Figure 4).

**Figure 7.**
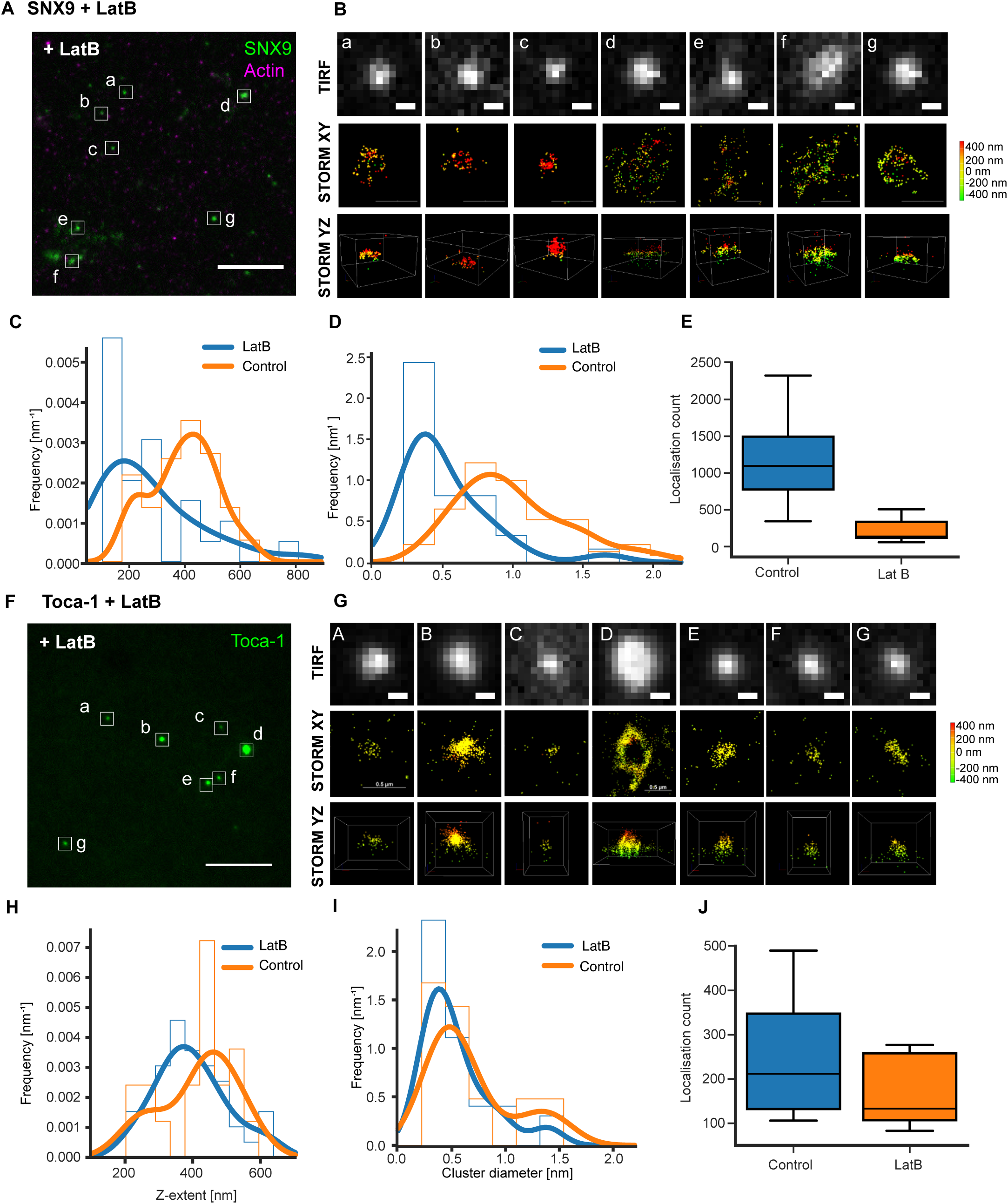
3D STORM imaging shows tubular, plaque-like and hybrid morphologies are retained yet reduced with actin monomer sequestering drug Latrunculin B, (A) Representative microscopy image showing SNX9 (green) by TIRF and actin by widefield (magenta). Scale bar 10 µm. (B) Closeup and STORM images of indicated FLS showing tubular, plaque-like and hybrid morphologies. The first row is the TIRF image with no discernable structural features. The second and third rows represent lateral and axial view of the 3D STORM reconstructions. Depth in the z-plane is colour coded from red (+400 nm) to green (-400 nm). Scale bar 0.5 µm. (C-E) LatB reduces the z-extent and cluster diameter as well as number of localizations. (F) Representative microscopy image showing TOCA-1 (green) by TIRF and actin by widefield (magenta). Scale bar 10 µm. (G) Closeup and STORM images of indicated FLS showing tubular, plaque-like and hybrid morphologies. The first row is the TIRF image with no discernable structural features. The second and third rows represent lateral and axial view of the 3D STORM reconstructions. Depth in the z-plane is colour coded from red (+400 nm) to green (-400 nm). Scale bar 0.5 µm. (H-J) LatB has little effect on TOCA-1 z-extent and cluster diameter or the number of localizations.

### Both branched actin comets and unbranched parallel actin bundles can arise downstream of SNX9 activity

Given the role for two Arp2/3 complex activators at filopodia tips (TOCA-1 and SNX9) as well as the tubular/endocytic types of morphologies we saw in STORM, we sought to verify the previous electron microscopy result that FLS are comprised of elongated and bundled actin filaments (Lee *et al*., 2010). FLS have previously been visualized by electron microscopy using phalloidin stabilization and formaldehyde fixation followed by negative stain, that showed the actin filaments tightly apposed and where they collapse and fray slightly, long, unbranched actin filaments are revealed. This agrees with the presence of fascin through the FLS shaft (Lee *et al*., 2010). With the recent progress in cryo-electron tomography (cryo-ET) methods, we took as less invasive a protocol as possible. To preserve FLS in their native form, we simply scratched the surface of the glass after 25 minutes of FLS growth on the supported bilayer (Figure 8A) and pipetted them onto EM grids. No branched actin comets were visible in any of our FLS tomograms. The resulting tomograms verify the bundled architecture (Figure 8B-C). To look at bundling more closely, we examined the FLS in transverse sections, where we saw areas with the characteristic hexagonal packing of fascin-bundled filaments (Figure 8B, insets a-b and Figure 8C inset a). Measurement of the inter-filament distances confirmed a ∼12 nm distance, consistent with measurements for fascin-mediated bundling *in vitro* (Yang *et al.,* 2013) and in filopodia (Hylton *et al.,* 2022; Atherton *et al.,* 2022) (Figure 8D).

**Figure 8.**
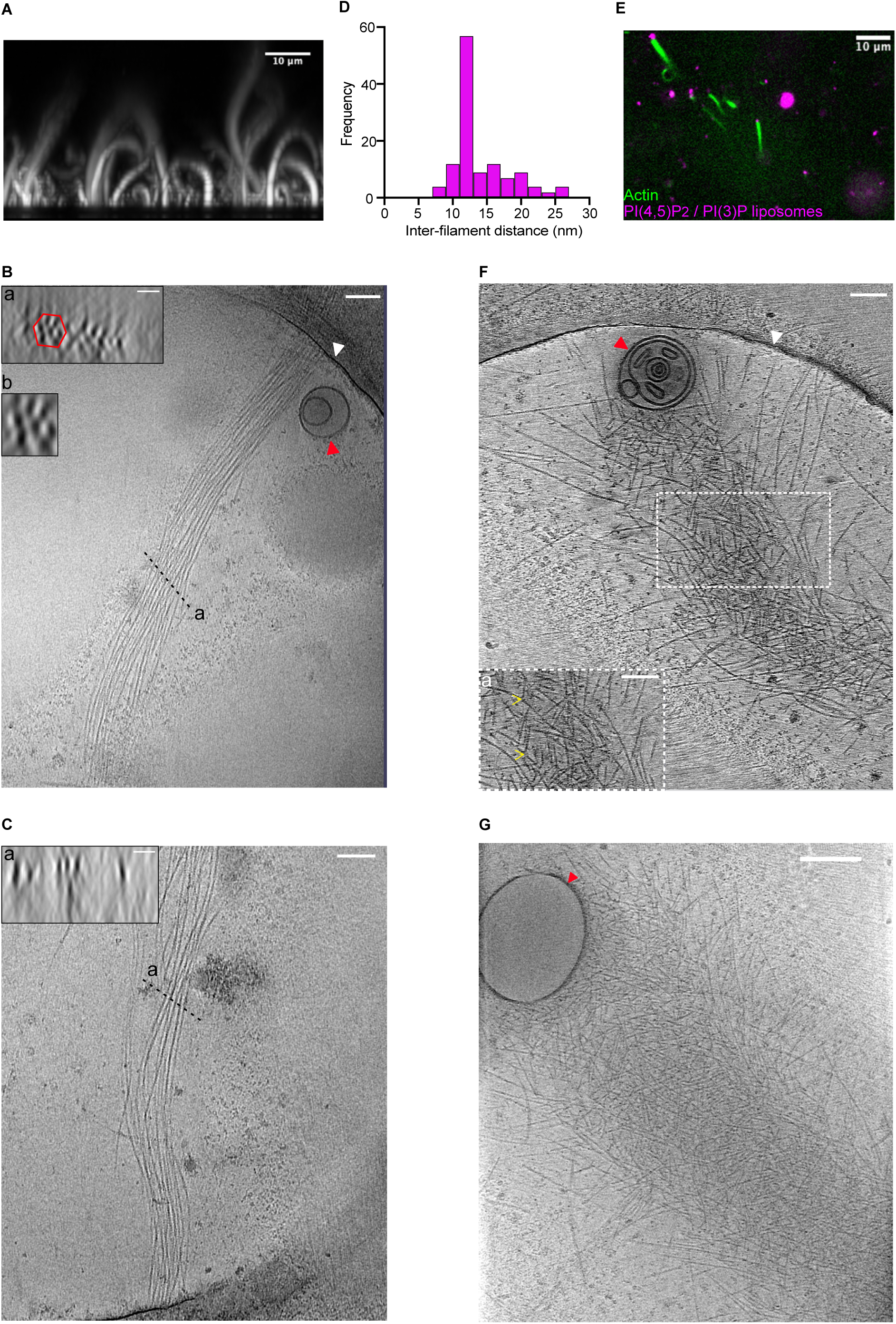
FLS grown from supported lipid bilayers have bundled and unbranched actin filament organization whereas comets grown from PI(4,5)P2/PI(3)P liposomes are highly branched. (A) Representative fluorescence image of FLS (B) Tomogram of an FLS, inset shows (a) zoom in of transverse section with actin filaments (b) showing hexagonal packing (scale bar = 25 nm) Red arrowhead indicates a liposome formed from membrane scratching during sample application on grid. White arrowhead indicates the edge of the grid hole. (C) Second example FLS. (D) Histogram of inter-filament distances (n = 118 across 4 tomograms) (E) Representative fluorescence image of actin comets coming from PI(4,5)P2/PI(3)P liposomes. Scale bar = 10 µm (F, G) Tomograms of representative comets, inset a shows closeup of branched filament organization, yellow arrowheads indicate branch points. Scale bars are 100 nm.

Using giant unilamellar vesicles and liposomes and the same type of frog egg extracts, we have previously attributed the effect of PI(3)P in stimulating actin via SNX9 and the Arp2/3 complex to an endocytic role and had assumed the actin was branched (Daste *et al*., 2017). We realised that instead the PI(3)P might be stimulating actin polymerization sufficiently to recruit elongation and bundling proteins and the previous actin we had attributed to comet tails could actually be long, unbranched and filopodia-like (Daste *et al*., 2017).

To test the actin architecture from PI(4,5)P2/PI(3)P containing small liposomes we previously used, that depend on SNX9 for their actin activity, we similarly applied them to EM grids and carried out cryo-ET (Figure 8E-G). In stark contrast to FLS, this actin is highly branched, with the branching maintained throughout the length of the network that originates from the liposome surface (Figure 8F-G, inset a shows a closeup of the branching indicated with yellow arrows). The branched nature of the filaments indeed agrees with expectations of endocytic actin observed in cells (Collins *et al*., 2011).

Our observations with SNX9 agree with the current two-phase model of FLS growth, with an Arp2/3-complex dependent nucleation and clustering phase followed by elongation and bundling by elongators and fascin, in line with the convergent elongation model of filopodia formation. Our data shows that the presence of long, bundled actin filaments is dependent on a unique feature of FLS growing from a PI(4,5)P2 supported bilayer. This actin morphology is not recapitulated by liposomes, despite the essential contribution of SNX9 to both, and presence of PI(3)P and curved SNX9/TOCA-1 intermediates at FLS (Figure 6).

## Discussion

We have shown that while half maximal binding concentrations of SNX9 for different phosphoinositide-containing liposomes is similar, the coincidence of PI(4,5)P2/PI(3)P allows more SNX9 to bind the liposome surface. This suggests that different multimeric arrangements of SNX9 are attained with different membrane composition. The difference in SNX9 binding to different membrane compositions is particularly acute at low SNX9 concentrations, which is the situation in cells, where SNX9 is present at approximately 120 nM (Wühr *et al*., 2014). We have shown that SNX9 recruitment to sites of actin polymerization is driven by lipids, its SH3 domain interaction partners and actin itself. This suggests that there is a cooperative network of membrane and actin regulators that centres around SNX9 organization. At FLS where SNX9 is involved in assembling actin, it can be present in either tubular or plaque-like morphologies, and we show evidence that actin and other factors in the extracts collaborate to bring about membrane deformation. SNX9 recruitment is reduced more than TOCA-1 in the presence of LatB, in agreement with previous data that TOCA-1 is an early protein to arrive at sites of FLS formation and its early role in nucleating filopodia themselves (Dobramysl *et al*., 2021; Blake *et al*., 2024).

Our work cannot distinguish whether the force provided by actin or BAR domains is the critical factor in membrane deformation. Both SNX9 and TOCA-1 localize to approximately 500 nm diameter deformations but can also localize to flat surfaces. The ability of membrane-localized actin to deform the membrane in either direction has been seen in purified reconstituted systems (Gat *et al*., 2020). Actin-generated BAR domain tubules emerge from native cell membranes in the presence of extracts, although here receptors and clathrin are also present that are absent in FLS (Wu *et al*., 2010). The assembly of TOCA-1 ortholog CIP4 oligomers on flat membranes has previously been observed in temperature cycles during clustering of the protein on membranes for cryoEM, with a specific mutation being identified (Frost *et al*., 2008).

We confirmed that FLS forming from PI(4,5)P2-containing supported lipid bilayers are comprised of long, parallel actin filaments, which may be due to TOCA-1 recruiting actin filament elongating proteins Enabled, Vasodilator-stimulated protein and Diaphanous homologs to the membrane (Dobramysl *et al*., 2021; Blake *et al*., 2024). Our observations agree with the previous two step (clustering-outgrowth) model of filopodia-like structure formation, with Arp2/3 complex-dependent actin nucleation as a limiting factor needed for recruitment of actin filament elongating and bundling proteins. The comparison of the actin superstructures by light microscopy and cryo-ET highlights the inability of light microscopy to distinguish actin bundling status.

Both PI(4,5)P2 and PI(3)P are present at both the supported lipid bilayers (by addition and conversion from the extracts) and in the small liposomes (by addition) and in both cases there is a SNX9 dependent activation of the Arp2/3 complex. It is an interesting open question how the ultimate difference in actin morphology arises.

The lack of effect of reducing SNX9 levels on filopodia formation in cells and lack of phenotype of the SNX9 knockout mouse (Liu *et al*., 2016) suggests that the initial SNX9-generated actin filaments formed that are the pre-requisite for growth in FLS can be provided by other molecular means in cells. Nonetheless we would predict that when the appropriate membrane signalling environment is present of PI(4,5)P2 and PI(3)P together with SH3 domain interaction partners, that SNX9 is a powerful co-factor in actin nucleation. We hypothesize that this process is a key consideration in the disease settings where SNX9 is involved.

## Materials and methods

### Lipids and protonation

All lipids used were purchased from Avanti Polar Lipids and were natural brain tissue for PC, PS and PI(4,5)P_2_ and synthetic for PI(3,4)P_2_ and PI(3)P. Lipids used were: 25 mg Brain PC (840053C), 25 mg brain PS (840032C), 1 mg brain PI (4,5)P_2_ (840046X-1 mg), 500 μg 18:1 PI(3)P (850150P-500UG), 100 or 500 μg 18:1 PI(3,4)P_2_ (850153P) and 18:1 Liss Rhodamine PE (810150C). Synthetic PI(3,4)P_2_ and PI(3)P powders were resuspended in CHCl_3_ directly in a glass vial, dried under a stream of nitrogen and dessicated for 1 h. Lipids were rehydrated with a 2 CHCl_3_: 1 MeOH: 0.01N HCl, incubated for 15 minutes, dried under nitrogen, followed by another 1 h dessication step. The resultant lipid film was resuspended in 3 CHCl_3_: 1 MeOH, dried under nitrogen, followed by another resuspension in CHCl_3_ and drying, and finally, a resuspension in CHCl_3_ at 0.1 or 0.5 mg/mL. Protonated lipids were stored at -80°C.

### Preparation of 100 nm liposomes

Prior to liposome formation, lipid aliquots stored in -80°C were allowed to equilibrate to room temperature. Following this, lipids were mixed to give the various to give a final concentration of 2 mM lipid. These were prepared by mixing lipid CHCl_3_/ MeOH stock in an appropriate quantity, drying under nitrogen followed by a 1.5 h desiccation. Lipids were rehydrated in either 1x XB (*Xenopus* extracts buffer: 100 mM KCl, 100 nM CaCl2, 1 mM MgCl2, 50 mM sucrose, and 10 mM K-Hepes, pH 7.4) to visualise actin comet formation on liposomes or HEPES-buffered saline pH 7.4 for BLI experiments. Rehydrated liposomes were subject to a freeze-thaw cycle 8 times, after which they were probe sonicated for 1 minute (2 s on 5 s off) at 50% amplitude with a 8 mm probe tip. To achieve 100 nm sized liposomes, probe sonicated liposomes were extruded 11 times, sequentially through 800 nm, 300 nm and 100 nm polycarbonate filter (Mini extruder; Avanti Polar lipids).

### FLS assay

The FLS assay was performed as described (Walrant *et al*., 2015). For a 50 μL FLS mix: 8.4 μL HSS (High Speed Supernatant ∼25 mg/ml), 2.5 μL 20x Energy mix (150 mM phosphocreatine, 20 mM Mg-ATP, 20 mM MgCl2), 0.5 μL 200 mM DTT, 1 μL Actin (unlabelled) 0.3 mg/mL in G-buffer (unlabelled), 0.6 μL 10x XB and 37 μL 1x XB. One day prior to the experiment either Atto- or Alexa-Fluor rabbit skeletal fluorescently labelled (Cytoskeleton) lyophilised aliquots were resuspended G-buffer (5 mM Tris-HCl pH 8.0, 0.2 mM CaCl2, 0.2 mM Na-ATP) overnight at 4°C. This gave a final working concentration of 0.3 mg/mL. Following this, the actin was spun down using a TLA-100 rotor (Beckman Coulter) at 55,000 rpm for 30 minutes at 4°C. The supernatant was snap frozen in aliquots for later FLS experiments. For all experiments all components were stored on ice, once mixed, the FLS mix (including all fluorescent proteins and chemical inhibitors) was pre-incubated for 10 minutes at room temperature, before adding to supported lipid bilayers. Wortmannin (Merck W-1628) was added at a 2 µM final concentration, while Dynole 31-2 or 34-2 (abcam ab120474) were both added at a 10 µM concentration. The DMSO vehicle control was added at an appropriate dilution to match the volume of inhibitor added. SNAP tagged SNX9 or dynamin 2 were chemically labelled with SNAP-Surface Alexa Fluor 488 or 647 (New England Biolabs S9129 / S9136) and included in the FLS mix at 30 nM or 25 nM final concentration respectively. mCh-2xFYVE and eGFP-PH2x(TAPP) purified probes were added at 1 µg/ml final concentration. All 3D reconstructions were made in Fiji (ImageJ). For cryo-ET of FLS, the assay was conducted at the Department of Biochemistry, University of Cambridge to minimise disruption of formed FLS structures. At 25 minutes, ∼40 µL of excess assay volume was removed. The glass coverslip was scratched with a pipette and applied to the grid before vitrification using a Vitrobot.

This research has been regulated under the Animals (Scientific Procedures) Act 1986 Amendment Regulations 2012 after ethical review by the University of Cambridge Animal Welfare and Ethical Review Body and covered by Home Office Project Licence PP1038769 (licence holder: J.L. Gallop) and Home Office Personal licences held by H. Fisher, T. Jones-Green, and J. Mason.

### Immunodepletion of SNX9 from extracts and wild-type SNX9, SNX9-PX mutant and SNX9-BAR mutant add-backs

Three rounds of immunodepletion and three rounds of mock depletion were performed using 25 µL protein A beads against 100 µL of *Xenopus laevis* HSS egg extracts. The beads were incubated with 4517 rabbit anti-SNX9 serum for 1 h at room temperature on a rotator. Beads were washed 8 times at room temperature, using the magnet to separate beads from wash, as follows - 2 x 500 µL PBST, 1x 500 µL PBST + 50 µL 5 M NaCl, 1x 500 µL PBST and 3x 500 µL 1x XB. Beads were then resuspended in 100 µL 1 x XB per 25 µL of beads. Anti-SNX9 bound beads were divided into three 25 µL samples. At 4°C. XB was removed and 100 µL of *Xenopus* egg extracts were added to one of the anti-SNX9 samples and 100 µL to the mock beads. The beads were incubated for 15- minutes at 4°C on a rotator (other two anti-SNX9 beads remained on the magnet at 4°C (until required). After the first round of depletion, beads were separated using a magnetic rack. Depleted extracts were removed and added to the next aliquot of beads. This was repeated 3 times with beads incubated for 15 minutes at 4°C on a rotator. Mock depletion beads were treated the same as the depletion beads but were incubated with pre-bleed of the rabbit anti-SNX9 serum.

### TIRF and confocal imaging of FLS

Imaging of all FLS experiments was performed on a custom combined spinning disk/total internal reflection (TIRF) fluorescence microscope supplied by Cairn research. The system was based on an Eclipse Ti-E inverted microscope (Nikon), fitted with an X-Light Nipkow spinning disk (Core Research for Evolutional Science and Technology), an iLas2 illuminator (Roper Scientific), a Spectra X LED illuminator (Lumencor), and a 250-µm piezo-driven Z-stage/controller (NanoScanZ, Prior). Images were collected at room temperature through a 100× 1.49 NA oil objective using a Photometrics Evolve Delta EMCCD camera in 16-bit depth using Metamorph software (Version 7.8.2.0, Molecular Devices). Alexa Fluor 488 and GFP samples were visualized using 470/40 excitation and 525/50 emission filters, Alexa Fluor 568 and mCherry samples with 560/25 excitation and 585/50 emission filters, and Alexa Fluor 647 samples with 628/40 excitation and 700/75 emission filters. For FLS assays, specific proteins of interest at the FLS tip (at the membrane) were imaged in TIRF microscopy, in conjunction with a confocal z-stack of the actin structure. Images were processed in FIJI (ImageJ; Schindelin et al., 2012).

### Quantifications of FLS parameters

The properties of FLS were measured and segmented using an ImageJ plug-in ‘FLS-Ace 2.0’. This plug-in was developed in collaboration between Richard Butler, Iris Jarsch and Ulrich Dobramysl and is available on Github. This software allows for the batch processing of large datasets to extract FLS parameters. Briefly, the FLS were segmented by a 2D Difference of Gaussians filter followed by the thresholding of every z-slice. The FLS were then tracked from the base of the structure to the tip. Additionally, other aspects such as base area, length and straightness were quantified. Threshold parameters were applied to identify ‘real FLS’. These were defined by a circularity above 0.5, tip area falling below 20 μm^2^ and a minimum length of 3 μm.

### Expi293FTM cell line expression and protein purification of full-length SNX9 and mutants

*X. laevis* SNX9 (accession no. <u>BC077183</u>) and mutants have been described previously (Daste *et al*., 2017). Expi293FTM were maintained in FreeStyle 293 Expression Medium (Thermo Fisher Scientific) supplemented with 100 μg/ml penicillin/100 U/ml streptomycin (Gibco). The cell line was grown at 37°C in a humified incubator shaking at 150 RPM with 8% CO_2_. pCS 6xHis-SNAP SNX9 constructs were transfected into 293F cells using 293fectin (Thermo Fisher Scientific) and were harvested 48 hours post transfection, according to the manufacturer’s instructions. Cells were harvested by centrifugation and resuspended in buffer containing 400 mM KCl, 20 mM K-Hepes, pH 7.4, 2 mM 2-*β*mercaptoethanol, and EDTA-free cOmplete protease inhibitor tablets and following snap-freezing using liquid nitrogen. Protein pellets were stored at -20°C.

Frozen pellets were thawed on ice until lysate has completely thawed. These pellets were lysed using probe sonication using 5 cycles (2 seconds on and 5 seconds off) of probe sonication at 70% amplitude. Lysates were spun at 45,000 RPM for 40 minutes (Optima MAX-XPN ultracentrifuge) followed by affinity purification. Ni-NTA beads (Qiagen) were used for 6xHis-tagged protein. 6xHis-tagged proteins were eluted from Ni-NTA beads in a stepwise manner with increasing concentrations of imidazole: 50-300 mM in buffer A: 150 mM NaCl, 20 mM HEPES pH 7.4 and 2 mM *β*-mercaptoethanol. Fractions from all elutions were pooled and spin concentrated (Merck 20, 000 MWCO) to 2 ml prior to gel filtration by flow pressure liquid chromatography using an AKTA pure. using the S200 16/600 column (Cytiva). Eluents were subjected to SDS-PAGE analysis to identify pure fractions before being aliquoted with 10% glycerol, snap-frozen and stored at -80°C.

### SDS-PAGE, Coomassie staining of protein purification and western-blot analysis of immunodepletions

30 μL of affinity purified, gel-filtered eluate, or 1:10 dilution of lysate/ depleted lysate with 1x Sample buffer was boiled at 95°C for 2-5 minutes. Samples were loaded on NuPAGE bis/tris 4-20% precast gels (BioRad). Electrophoresis was done for 40 minutes at 185 V. Polyacrylamide gels for Coomassie stained were incubated for 30-minutes or overnight with Coomassie stain. After incubation the gels were destained overnight with distilled water. For western blot analysis, 2 µg of extracts (either depleted, mock or untreated) were diluted with 1x sample buffer, 10 µL of which was used for gel0electrophoresis. After run, polyacrylamide gels were transferred to a nitrocellulose membrane using iBlot 2 Dry Blotting system (for 7 minutes at 20V). After transfer, membrane was blocked with 5% milk in TBST. Incubation with primary antibody (polyclonal anti-SNX9 antibody) diluted in 1% milk in TBST overnight at 4°C. This was followed by washing 3 times with PBST. Secondary antibody incubation was done for 1 h at room temperature followed by washing 3 times in TBST. Membrane was visualized using LI-COR Odyssey clx.

### Biolayer Interferometry (BLI)

BLI experiments were conducted on Octet RED96 (Forté Bio) for the analysis of SNX9 binding to phosphatidylinositol lipids. All experiments were conducted using Aminopropylsilane (APS) biosensors (Sartorius). APS sensors were washed in baseline buffer: HBS-N buffer supplemented with 2 mM DTT. Liposomes were immobilised onto the APS surface for 1500 s, after which the sensor was washed with baseline buffer. The surface was then blocked with 0.5% BSA for 600 s, succeeded by another wash step. The baseline was allowed to stabilise before studying protein binding. The probe was moved to well with protein, allowing for protein to bind to liposomes (association step) which continued for 2000 s, after which the probe was moved into a buffer only well for 1500 s to promote dissociation of proteins. All binding curves were corrected with a PC/PS only control and were then corrected to the baseline preceding the association phase.

### Surface Plasmon Resonance

A manual SPR run was conducted using BIAcore T200 using a Series S L1 chip (Cytiva). Liposomes were individually immobilised (∼25 minutes) with a flow rate of 2 μL/ min. Post immobilisation, all channels in the chip were washed for 30 s with HEPES-buffer saline (HBS-N) pH 7.4, to remove unbound liposomes. SNX9 was subsequently injected over all 4 channels at 2 μL/ min for 10-minutes. Dissociation of SNX9 was allowed to occur over 30-minutes, following which, the final 30 s wash (2 µL/min) with HBS-N was conducted before, the dextran matrix was regenerated with a 2x 30 s 50 mM NaOH/ isopropanol (2 µL/ min). System and chip were washed as per manufacturer’s instructions.

### Actin comet formation on LUVs

This assay was conducted on a glass coverslip with the use of a silicone gasket. The glass in the resultant wells was passivated with 5% fatty acid free bovine serum albumin (Sigma) for 90 minutes. The wells were washed three times with 1x XB. Wells were left in 1x XB until they were ready for use. Liposomes for the actin formation on LUVs were filtered to 100 nm and comprised of 65% PC / 30% PS / 4% PI(4,5)P2/ 1% PI(3)P. For the assay, liposomes were diluted: 3 μL / 25 μL in 1x XB after which they were added to the well. These were allowed to settle for 10 minutes, after which a 25 μL ‘actin-mix’ was added including 2 mM DTT, 3.6 mg / mL *Xenopus laevis* egg extracts, and 1x Energy mix. Comets were allowed to form for 20 minutes and 4 µL of the sample was applied to the grid for vitrification.

### Vitrification of FLS and actin comets on LUVs

FLS or actin comets (both without fluorescently tagged actin) were applied to glow discharged R2/2 Cu 200 mesh grids (PELCO Easiglow discharge cleaning system; current: 25 mA, time: 60 seconds, polarity: negative, vacuum: 0.39mBar, vacuum stabilisation wait time: 10 seconds) and blotted for 3 seconds followed by immediate plunge freezing in liquid ethane with an FEI Vitrobot Mark IV (ThermoScientific) (95% humidity at 4°C). Grids were first screened in a Talos Arctica equipped with a Falcon 3EC detector and operating at 200 kV (at the Department of Biochemistry, University of Cambridge), and grids of good quality were chosen for cryo-ET data collection.

### Cryo-electron tomography and data processing

Cryo-ET acquisition was conducted at two separate facilities. For actin comets on LUVs, data collection was performed at the London Consortium for Cryo-EM (LonCEM) facility using a Titan Krios G3i microscope operating at 300 keV equipped with a Gatan K3 direct electron detector. Tilt series were collected in K3 super-resolution mode using a dose-symmetric protocol with a dose of 2.96 e^-^/tilt and a step size of 3° starting from 0° and progressing to ±60°. The defocus range was set between -2.5 µm and -5 µm, with a physical pixel size of 3.33 Å. For FLS, data were collected at the Birkbeck facility on a Titan Krios microscope operating at 300 keV with a K3 detector in super-resolution mode. Tilt series were collected using a dose-symmetric tilt scheme, starting from 0° and incrementing by 3° up to ±60°. A physical pixel size of 3.4 Å was used, and the defocus range was set between -3.0 µm and -7.0 µm. Fiducial-less tilt series alignment was performed with AreTomo v1.4.3 (Zheng *et al*., 2022) followed by Wiener-filtered tomogram reconstruction in either RELION-4 (with binning factor of 4, final pixel size of 13.3 Å) (Kimanius *et al.,* 2021) or RELION-5 (with a binning factor of 6, final pixel size of 20.4 Å) (Burt *et al.,* 2024) for the actin comets or FLS datasets respectively.

Representative images were created in IMOD software (Kremer *et al*., 1996). To determine the inter-filament distances, the tomogram was rotated in the slicer window of IMOD to identify a section with a cross-section through the actin bundle. One actin filament was centered in the field of view as a reference point and the centres of neighbouring filaments identified by clicking ‘Q’ on keyboard to read out the distance. These distances were reported in nm and plotted using GraphPad (Prism).

### Preparation of FLS for super-resolution imaging using direct Stochastic Optical Reconstruction Microscopy (dSTORM)

FLS were stabilized by adding 20 µM unlabeled phalloidin (Molecular Probes, no. P3457) for 5 min. FLS were carefully washed with XB buffer twice to remove any loose actin floating in solution and fixed in 4% formaldehyde for at least 60 min. Fixed FLS were washed three times with XB to remove the formaldehyde. Stock solutions of STORM buffer components were mixed together at a ratio of 30 µL, 500 µL, and 60 µL: enzyme mix [20 µg/ml catalase (Sigma), 20 mM Tris-HCl (pH 7.5), 1 mg/ml glucose oxidase (Sigma)], glucose solution [100 mg/ml D-Glucose (Sigma) in ddH2O], and reducing agent [1M cysteamine mercaptoethylamine (MEA, Sigma) in ddH2O] prior to adding to FLS and imaging. 20 µM of Latrunculin B in DMSO (Lat B; Merck, no. 428020) was incubated in the FLS reaction mixture for 5 min, before the mix was added to the lipid bilayer. The brightness/contrast range of the protein Total Internal Reflection Fluorescence (TIRF) images were adjusted uniformly in ImageJ/Fiji (National Institutes of Health; Schindelin et al., 2012), and the local maxima was identified using the *Find Maxima* function.

### dSTORM data acquisition

A Nikon STORM (N-STORM) system as used with an Agilent laser bed with 405 nm, 488 nm, 561 nm, and 647 nm lasers. The emitted light was collected through a CPI Plan Apo 100X 1.49 NA objective, filtered by a bandpass filter (for Alexa Fluor 525/50; for Alexa Fluor 568: 600/50; for Alexa Fluor 647: 705/72; for mEos and multi-color imaging: NSTORM QUAD 405/488/561/647) and imaged onto an iXon Ultra 897 EMCCD camera (Andor). The setup was controlled by Nikon NIS-Elements AR software version 4.50 with N-STORM module. A reference TIRF image of the protein of interest and actin was acquired using a widefield Z stack. The cylindrical lens and condenser lens were inserted into the light path, and the focus was . Movies of typically 30,000 frames (size 256×256 pixels, equivalent to 39.9×39.9 µm as 1 pixel = 0.16 µm) were acquired using 647 nm illumination at 2-3 kW/cm^2^ at 19 ms exposure times, once the fluorophores had transitioned into a sparse blinking pattern. For samples of low fluorescence intensity, 405 nm laser power with an automated feedback loop was adjusted to keep a constant number of localizations per frame.

### dSTORM analysis

Coordinates of localization points were obtained and used to create 3D reconstructions using the N-STORM module in NIS-Elements AR software. Sample drift and chromatic aberration was corrected for using the 3D bead calibration sample. To exclude camera noise, the peak detection threshold was set to 4,000-5,000 gray levels. FLS tips which showed co-localization of actin and the labeled protein of interest by TIRF and a length of actin growing at least 2 µm in the z-axis were selected. Histograms of localization positions in Z were also obtained and normalized to the mean Z positions to compare over multiple FLS. Substructures of SNX9 and TOCA-1 were classified into plaques, tubules or hybrid structures according to how their 3D geometries appeared. Tubules were identified as circular objects with apparent Z height rising above the membrane, most often with more localizations on its periphery than in its core. Plaques were identified as flatter structures with ridged edges. Hybrid structures were identified as intermediate structures that displayed both properties within the same tip regions, for example a tubule-like structure seemingly emerging from a part of the plaque-like platform. The Z-extent and diameter measurements were also measured for each of these subgroups, to quantify the differences in their properties.

## Supplementary Figure Legends

**Sup. Figure 1.**
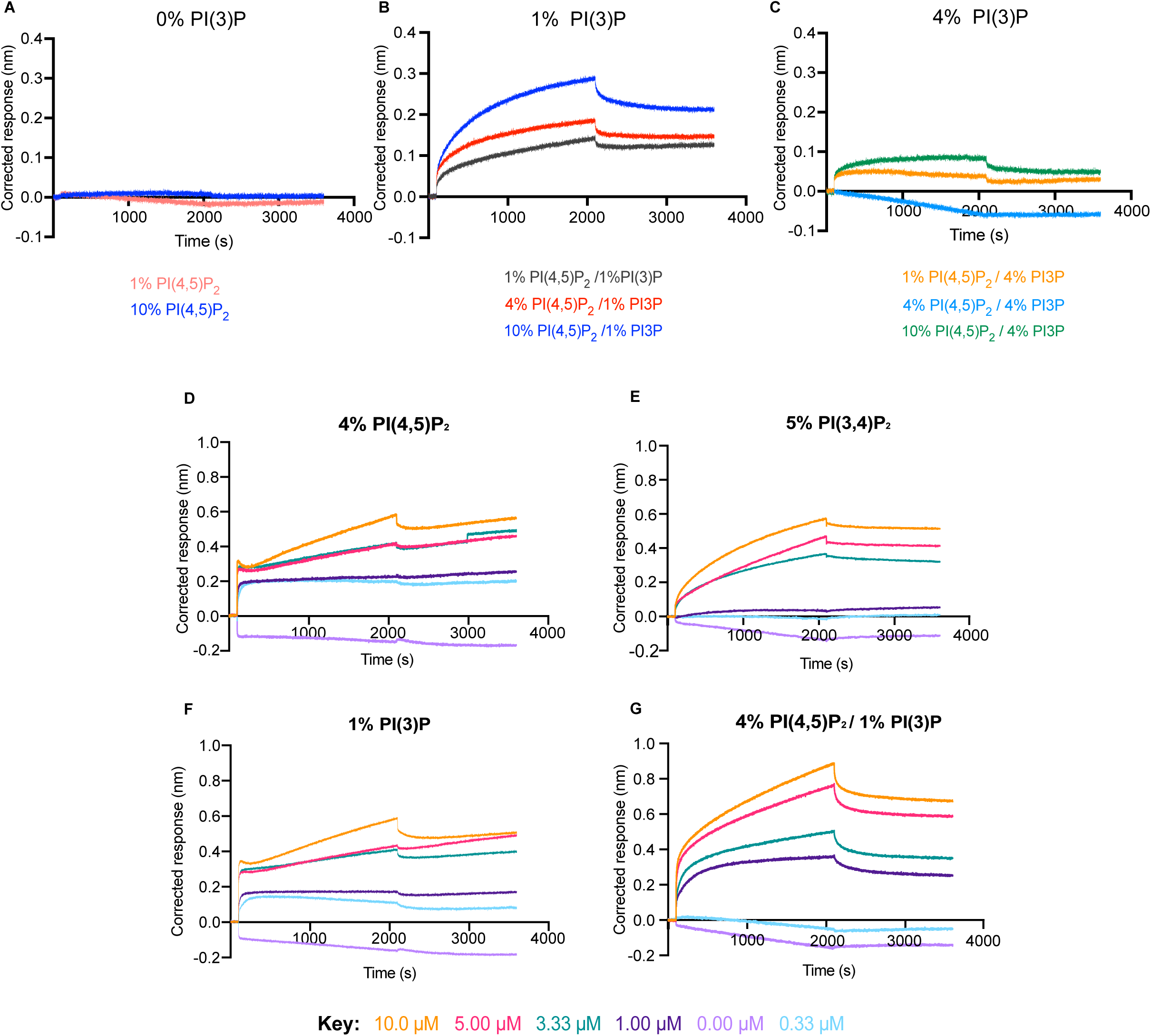
BLI traces with different lipid compositions corresponding to Figure 1D-E. All traces show the association and dissociation phases only, background subtracted to 70% PC / 30% PS liposomes. (A) At 1 µM SNX9 there is no additional binding with 1 or 10% PI(4,5)P2. (B) Increased binding with increased PI(4,5)P2 with 1% PI(3)P at 1 µM SNX9. (C) Increasing PI(3)P further is inhibitory. (D-G) Distinct binding behaviour with the individual and combined lipids indicated at SNX9 concentrations ranging from 0.33 µM to 10 µM.

**Sup. Figure 2.**
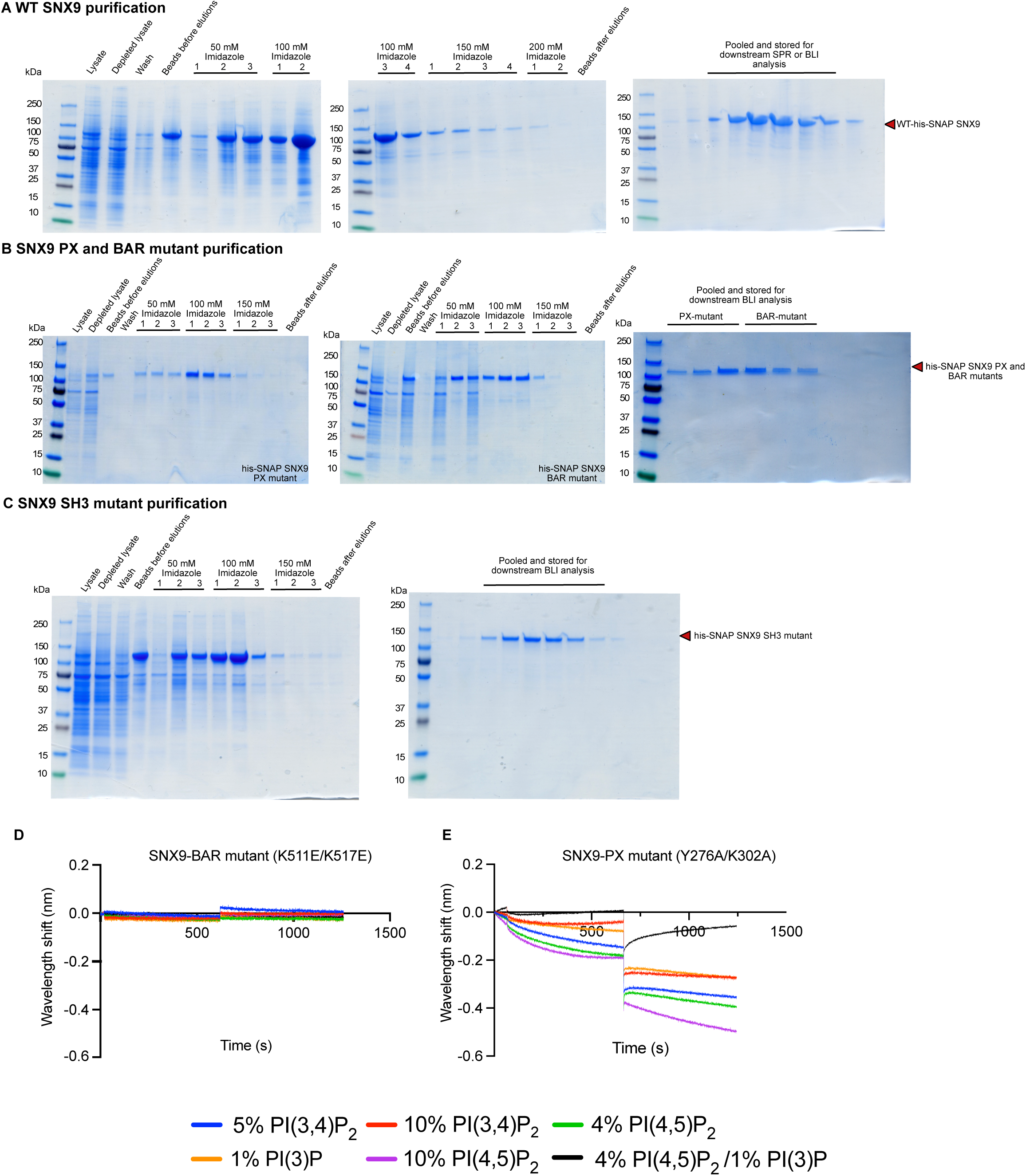
The SNX9 BAR and PX domain mutants do not bind effectively to any of the lipid compositions. (A) Protein purifications of SNX9 (B) PX and BAR domain mutants (C) SH3 domain mutants. (D) BLI traces with full-length SNX9 K511E/K517E with indicated phosphoinositide compositions in 60-69% mol fraction PC and 30% mol fraction PS. (E) BLI traces with full length SNX9 Y276A/K302A show no binding.

**Sup. Figure 3.**
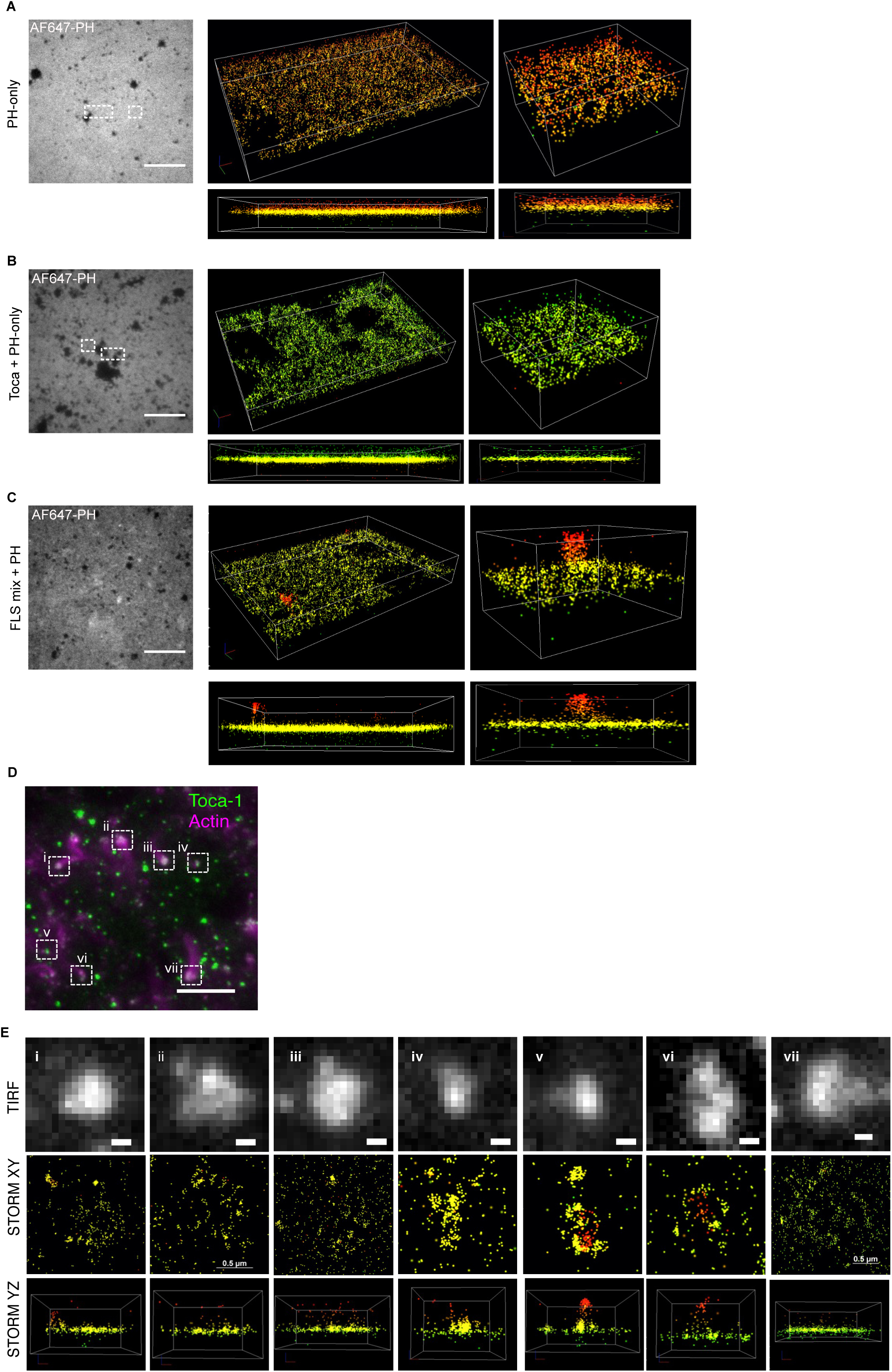
3D-STORM shows tubular, plaque-like and hybrid morphologies of TOCA-1 at FLS sites (A) Representative microscopy image showing PLCd-PH domain bound to the membrane and flat localizations by STORM. (B) Similarly flat overall binding with TOCA-1 plus PLCd PH domain (C) When extracts are added, some PH domain clustering and tubular morphology is seen. (D) TOCA-1 (green) by TIRF and actin by widefield (magenta). Scale bar 10 µm. (E) Closeup and STORM images of indicated FLS showing tubular, plaque-like and hybrid morphologies. The first row is the TIRF image with no discernable structural features. The second and third rows represent lateral and axial view of the 3D STORM reconstructions. Depth in the z-plane is colour coded from red (+400 nm) to green (-400 nm). Scale bar 0.5 µm.

**Sup. Figure 4.**
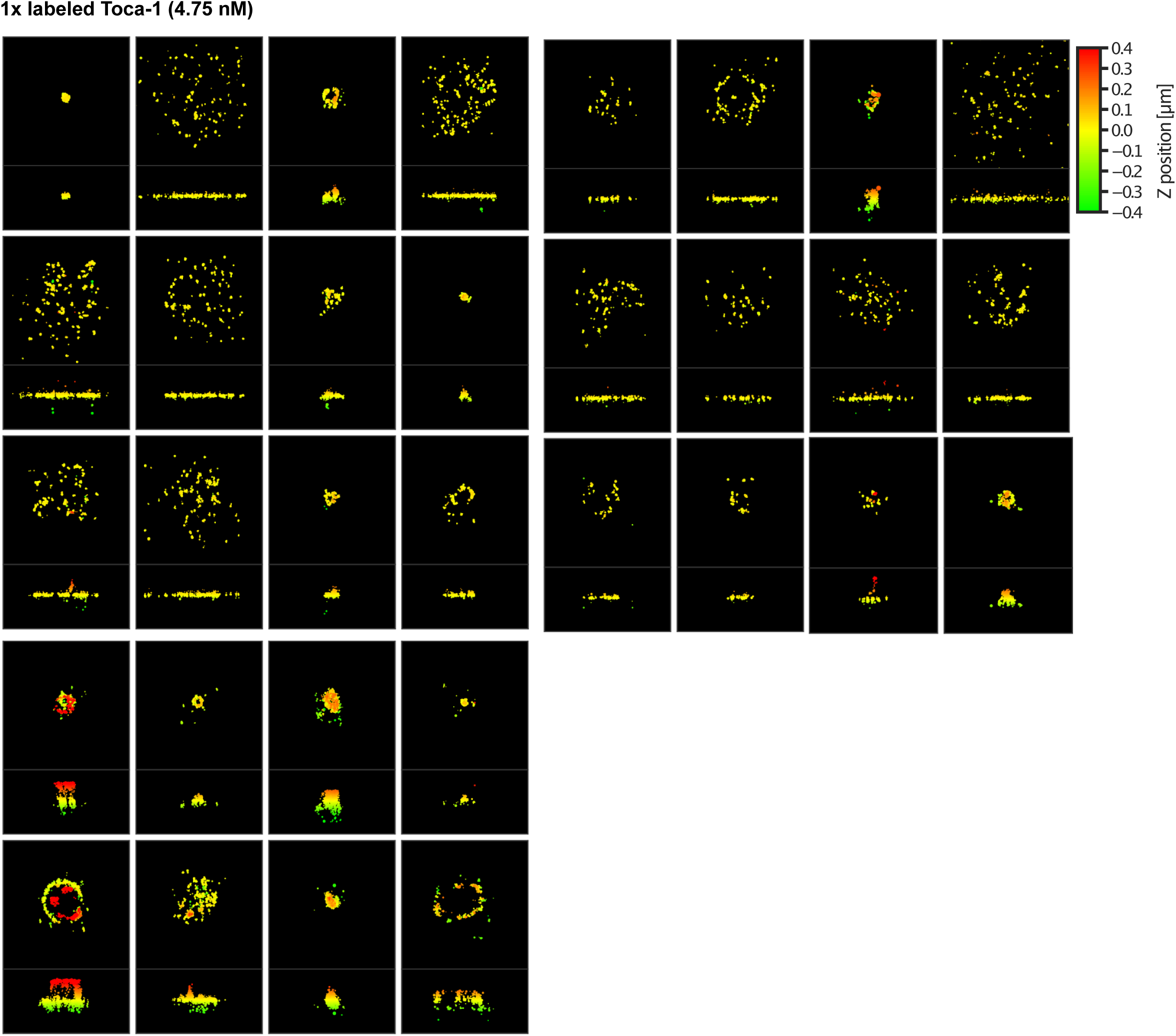
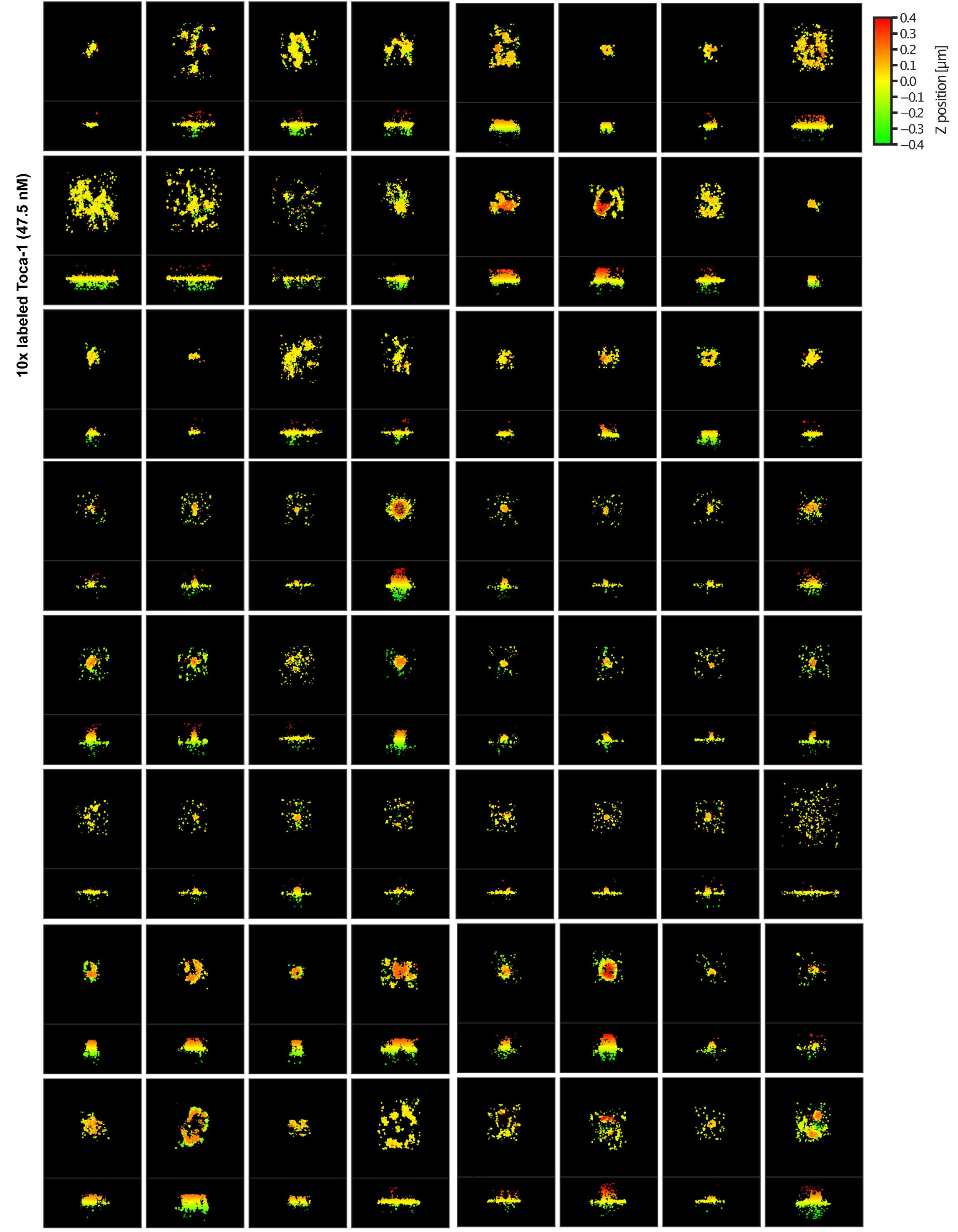
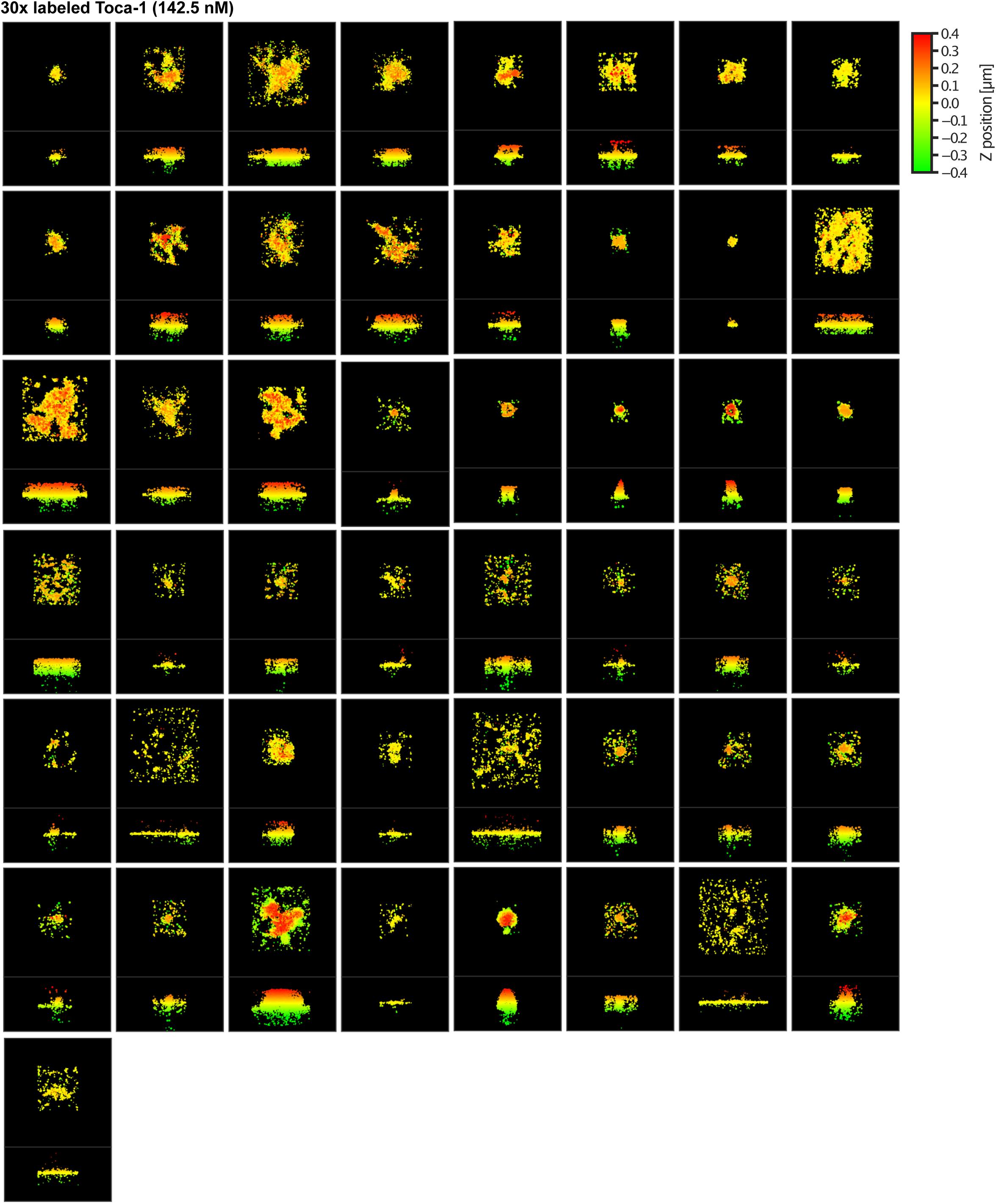
Gallery of TOCA-1 morphologies at FLS at increasing TOCA-1 concentrations showing that tubules and plaques are found similarly at all additional TOCA-1 concentrations.

**Sup. Figure 5.**
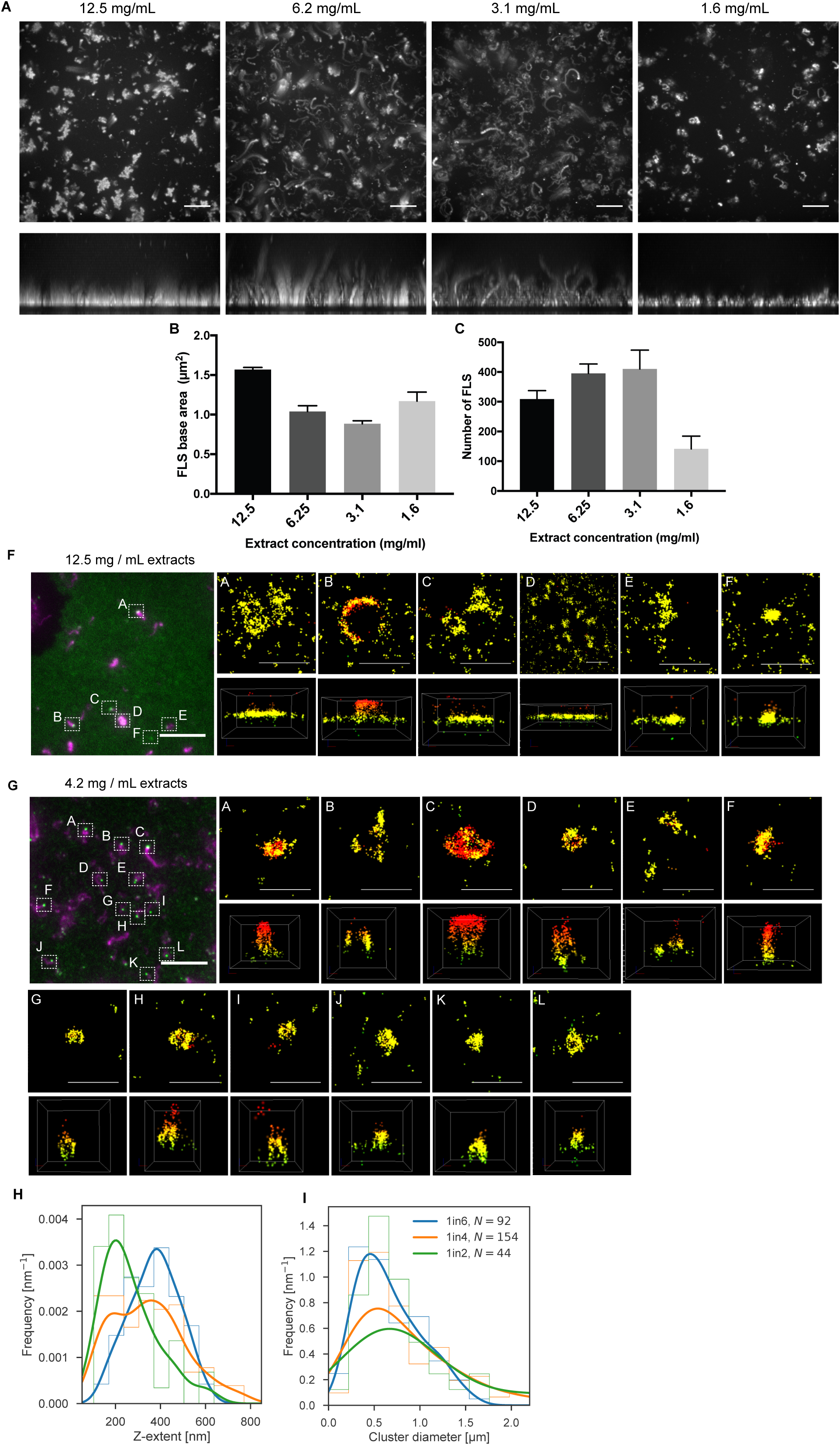
Gallery of TOCA-1 morphologies at FLS at increasing extract concentrations showing that more plaques are observed at higher concentrations and more tubules at lower concentrations. (A) Confocal micrographs of x-y and z views of FLS under different extract conditions. The FLS lengths (of actin) are longest low to intermediate concentrations and there are more formed. 1 in 6 = 3.1 mg/ml, 1 in 4 = 6.2 mg/ml, 1 in 2 = 12.4 mg/ml extract concentrations.

## Acknowledgments

We thank Astrid Walrant, Thomas Blake, Mary-Pia Jeyarajasingham and Laura Machesky for valuable discussion and reading the manuscript. We thank Mr Lee Cooper, Dr Steve W. Hardwick and Dr Dimitri Y. Chirgadze for assistance with data collection at the Cryo-EM Facility, Department of Biochemistry, University of Cambridge. The facility was established using funds from the Wellcome Trust (206171/Z/17/Z; 202905/Z/16/Z) and the University of Cambridge. Cryo-EM were also collected at LonCEM which is supported by Wellcome Grant no: 206175/Z/17/Z and partner institutes and at Institute of Structural and Molecular Biology EM facility at Birkbeck College, University of London with financial support from the Wellcome Trust (202679/Z/16/Z and 206166/Z/17/Z). We thank Katherine Stott and the Biophysics Facility in the Department of Biochemistry, University of Cambridge for access to SPR and BLI. This work was funded by a Wellcome Trust Senior Research Fellowship 219482/Z/19/Z, Wellcome Trust Research Career Development Fellowship WT095829MA and Issac Newton Trust equipment grant to JLG and a Funai Foundation studentship to HSK. For the purpose of open access, the author has applied a Creative Commons Attribution (CC BY) licence to any Author Accepted Manuscript version arising from this submission. The eGFP-PH2xTAPP construct was a kind gift from Volker Haucke and Michael Krauss.

## Notes

### Competing Interest Statement

The authors have declared no competing interest.

## References

Atherton, J, Stouffer, M, Francis, F, and Moores, CA (2022). Visualising the cytoskeletal machinery in neuronal growth cones using cryo-electron tomography. J Cell Sci 135, jcs259234.

Bendris, N, and Schmid, SL (2017). Endocytosis, Metastasis and Beyond: Multiple Facets of SNX9. Trends in Cell Biology 27, 189–200.

Bendris, N, Stearns, CJS, Reis, CR, Rodriguez-Canales, J, Liu, H, Witkiewicz, AW, and Schmid, SL (2016a). Sorting nexin 9 negatively regulates invadopodia formation and function in cancer cells. Journal of Cell Science 129, 2804–2816.

Bendris, N, Williams, KC, Reis, CR, Welf, ES, Chen, PH, Lemmers, B, Hahne, M, Leong, HS, and Schmid, SL (2016b). SNX9 promotes metastasis by enhancing cancer cell invasion via differential regulation of RhoGTPases. Molecular Biology of the Cell 27, 1409–1419.

Blake, TCA, Fox, HM, Urbančič, V, Ravishankar, R, Wolowczyk, A, Allgeyer, ES, Mason, J, Danuser, G, and Gallop, JL (2024). Filopodial protrusion driven by density-dependent Ena-TOCA-1 interactions. J Cell Sci 137, jcs261057.

Burt, A, Toader, B, Warshamanage, R, Kügelgen, A von, Pyle, E, Zivanov, J, Kimanius, D, Bharat, TAM, and Scheres, SHW (2024). An image processing pipeline for electron cryo-tomography in RELION-5. FEBS Open Bio.

Chandra, M, Chin, YK-Y, Mas, C, Feathers, JR, Paul, B, Datta, S, Chen, K-E, Jia, X, Yang, Z, Norwood, SJ, et al. (2019). Classification of the human phox homology (PX) domains based on their phosphoinositide binding specificities. Nat Commun 10, 1528.

Collins, A, Warrington, A, Taylor, KA, and Svitkina, T (2011). Structural Organization of the Actin Cytoskeleton at Sites of Clathrin-Mediated Endocytosis. Curr Biol 21, 1167– 1175.

Daste, F, Walrant, A, Holst, MR, Gadsby, JR, Mason, J, Lee, JE, Brook, D, Mettlen, M, Larsson, E, Lee, SF, et al. (2017). Control of actin polymerization via the coincidence of phosphoinositides and high membrane curvature. Journal of Cell Biology 216, 3745– 3765.

Dobramysl, U, Jarsch, IK, Inoue, Y, Shimo, H, Richier, B, Gadsby, JR, Mason, J, Szałapak, A, Ioannou, PS, Correia, GP, et al. (2021). Stochastic combinations of actin regulatory proteins are sufficient to drive filopodia formation. J Cell Biology 220, e202003052.

Dowler, S, Currie, RA, Campbell, DG, Kular, G, Deak, M, Downes, CP, and Alessi, DR (2000). Identification of pleckstrin-homology-domain-containing proteins with novel phosphoinositide-binding specificities. Biochem J 351, 19.

Ecker, M, Schregle, R, Kapoor-Kaushik, N, Rossatti, P, Betzler, VM, Kempe, D, Biro, M, Ariotti, N, Redpath, GM, and Rossy, J (2022). SNX9-induced membrane tubulation regulates CD28 cluster stability and signalling. ELife 11, e67550.

Ford, C, Nans, A, Boucrot, E, and Hayward, RD (2018). Chlamydia exploits filopodial capture and a macropinocytosis-like pathway for host cell entry. PLOS Pathogens 14, e1007051.

Frost, A, Perera, R, Roux, A, Spasov, K, Destaing, O, Egelman, EH, Camilli, PD, and Unger, VM (2008). Structural Basis of Membrane Invagination by F-BAR Domains. Cell 132, 807–817.

Gallop, JL, Walrant, A, Cantley, LC, and Kirschner, MW (2013). Phosphoinositides and membrane curvature switch the mode of actin polymerization via selective recruitment of toca-1 and Snx9. Proc National Acad Sci 110, 7193–7198.

Gat, S, Simon, C, Campillo, C, Bernheim-Groswasser, A, and Sykes, C (2020). Finger-like membrane protrusions are favored by heterogeneities in the actin network. Soft Matter 16, 7222–7230.

Gillooly, DJ, Morrow, IC, Lindsay, M, Gould, R, Bryant, NJ, Gaullier, J, Parton, RG, and Stenmark, H (2000). Localization of phosphatidylinositol 3-phosphate in yeast and mammalian cells. EMBO J 19, 4577–4588.

He, K, III, RM, Upadhyayula, S, Song, E, Dang, S, Capraro, BR, Wang, W, Skillern, W, Gaudin, R, Ma, M, et al. (2017). Dynamics of phosphoinositide conversion in clathrin-mediated endocytic traffic. Nature 552, 410–414.

Hicks, L, Liu, G, Ukken, FP, Lu, S, Bollinger, KE, O’Connor-Giles, K, and Gonsalvez, GB (2015). Depletion or over-expression of Sh3px1 results in dramatic changes in cell morphology. Biology Open 4, 1448–1461.

Hill, TA, Gordon, CP, McGeachie, AB, Venn-Brown, B, Odell, LR, Chau, N, Quan, A, Mariana, A, Sakoff, JA, Fabbro, MC (nee, et al. (2009). Inhibition of Dynamin Mediated Endocytosis by the Dynoles□Synthesis and Functional Activity of a Family of Indoles. J Med Chem 52, 3762–3773.

Hylton, RK, Heebner, JE, Grillo, MA, and Swulius, MT (2022). Cofilactin filaments regulate filopodial structure and dynamics in neuronal growth cones. Nat Commun 13, 2439.

Ish-Shalom, E, Meirow, Y, Sade-Feldman, M, Kanterman, J, Wang, L, Mizrahi, O, Klieger, Y, and Baniyash, M (2016). Impaired SNX9 Expression in Immune Cells during Chronic Inflammation: Prognostic and Diagnostic Implications. J Immunol 196, 156–167.

Jarsch, IK, Gadsby, J, Nuccitelli, A, Mason, J, Shimo, H, Pilloux, L, Marzook, Mulvey, CM, Dobramysl, U, Bradshaw, C Lilley, KS, Hayward, R, Vaughan, TJ, Dobson, CL, Gallop, J et al. (2020). A direct role for SNX9 in the biogenesis of filopodia. The Journal of Cell Biology 219.

Kimanius, D, Dong, L, Sharov, G, Nakane, T, and Scheres, SHW (2021). New tools for automated cryo-EM single-particle analysis in RELION-4.0. Biochem J 478, 4169–4185.

Kremer, JR, Mastronarde, DN, and McIntosh, JR (1996). Computer Visualization of Three-Dimensional Image Data Using IMOD. J Struct Biol 116, 71–76.

Lee, E, and Camilli, PD (2002). Dynamin at actin tails. Proc Natl Acad Sci 99, 161–166.

Lee, K, Gallop, JL, Rambani, K, and Kirschner, MW (2010). Self-Assembly of Filopodia-Like Structures on Supported Lipid Bilayers. Science 329, 1341–1345.

Liu, C, Zhai, X, Du, H, Cao, Y, Cao, H, Wang, Y, Yu, X, Gao, J, and Xu, Z (2016). Sorting nexin 9 (SNX9) is not essential for development and auditory function in mice. Oncotarget 7, 68921–68932.

Lo, W-T, Žagar, AV, Gerth, F, Lehmann, M, Puchkov, D, Krylova, O, Freund, C, Scapozza, L, Vadas, O, and Haucke, V (2017). A Coincidence Detection Mechanism Controls PX-BAR Domain-Mediated Endocytic Membrane Remodeling via an Allosteric Structural Switch. Dev Cell 43, 522–529.e4.

Lundmark, R, and Carlsson, SR (2003). Sorting Nexin 9 Participates in Clathrin-mediated Endocytosis through Interactions with the Core Components. Journal of Biological Chemistry 278, 46772–46781.

Park, J, Kim, Y, Lee, S, Park, JJ, Park, ZY, Sun, W, Kim, H, and Chang, S (2010). SNX18 shares a redundant role with SNX9 and modulates endocytic trafficking at the plasma membrane. Journal of Cell Science 123, 1742–1750.

Posor, Y, Eichhorn-Gruenig, M, Puchkov, D, Schöneberg, J, Ullrich, A, Lampe, A, Müller, R, Zarbakhsh, S, Gulluni, F, Hirsch, E, et al. (2013a). Spatiotemporal control of endocytosis by phosphatidylinositol-3,4-bisphosphate. Nature 2013 499:7457 499, 233–237.

Pylypenko, O, Lundmark, R, Rasmuson, E, Carlsson, SR, and Rak, A (2007). The PX-BAR membrane-remodeling unit of sorting nexin 9. EMBO J 26, 4788–4800.

Saengsawang, W, Mitok, K, Viesselmann, C, Pietila, L, Lumbard, DC, Corey, SJ, and Dent, EW (2012). The F-BAR Protein CIP4 Inhibits Neurite Formation by Producing Lamellipodial Protrusions. Curr Biol 22, 494–501.

Schöneberg, J, Lehmann, M, Ullrich, A, Posor, Y, Lo, WT, Lichtner, G, Schmoranzer, J, Haucke, V, and Noé, F (2017a). Lipid-mediated PX-BAR domain recruitment couples local membrane constriction to endocytic vesicle fission. Nature Communications 8, 1–17.

Shin, N, Ahn, N, Chang-Ileto, B, Park, J, Takei, K, Ahn, SG, Kim, SA, Paolo, GD, and Chang, S (2008). SNX9 regulates tubular invagination of the plasma membrane through interaction with actin cytoskeleton and dynamin 2. Journal of Cell Science 121, 1252– 1263.

Shukla, N, Kour, B, Sharma, D, Vijayvargiya, M, Sadasukhi, TC, Medicherla, KM, Malik, B, Bissa, B, Vuree, S, Lohiya, NK, et al. (2023). Towards Understanding the Key Signature Pathways Associated from Differentially Expressed Gene Analysis in an Indian Prostate Cancer Cohort. Diseases 11, 72.

Soulet, D, Y, M, L, and SL, S (2005). SNX9 regulates dynamin assembly and is required for efficient clathrin-mediated endocytosis. Molecular Biology of the Cell 16, 2058– 2067.

Tanigawa, K, Maekawa, M, Kiyoi, T, Nakayama, J, Kitazawa, R, Kitazawa, S, Semba, K, Taguchi, T, Akita, S, Yoshida, M, et al. (2019). SNX9 determines the surface levels of integrin β1 in vascular endothelial cells: Implication in poor prognosis of human colorectal cancers overexpressing SNX9. J Cell Physiol 234, 17280–17294.

Taylor, K, Pearson, M, Das, S, Sardell, J, Chocian, K, and Gardner, S (2023). Genetic risk factors for severe and fatigue dominant long COVID and commonalities with ME/CFS identified by combinatorial analysis. J Transl Med 21, 775.

Taylor, KL, Taylor, RJ, Richters, KE, Huynh, B, Carrington, J, McDermott, ME, Wilson, RL, and Dent, EW (2019). Opposing functions of F-BAR proteins in neuronal membrane protrusion, tubule formation, and neurite outgrowth. Life Sci Alliance 2, e201800288.

Taylor, MJ, Lampe, M, and Merrifield, CJ (2012). A Feedback Loop between Dynamin and Actin Recruitment during Clathrin-Mediated Endocytosis. PLoS Biol 10, e1001302.

Todkar, K, Chikhi, L, Desjardins, V, El-Mortada, F, Pépin, G, and Germain, M (2021). Selective packaging of mitochondrial proteins into extracellular vesicles prevents the release of mitochondrial DAMPs. Nat Commun 12, 1971.

Towers, CG, Wodetzki, DK, Thorburn, J, Smith, KR, Caino, MC, and Thorburn, A (2021). Mitochondrial-derived vesicles compensate for loss of LC3-mediated mitophagy. Dev Cell 56, 2029–2042.e5.

Trefny, MP, Kirchhammer, N, Maur, PA der, Natoli, M, Schmid, D, Germann, M, Rodriguez, LF, Herzig, P, Lötscher, J, Akrami, M, et al. (2023). Deletion of SNX9 alleviates CD8 T cell exhaustion for effective cellular cancer immunotherapy. Nat Commun 14, 86.

Walrant, A, Saxton, DS, Correia, GP, and Gallop, JL (2015). Triggering actin polymerization in Xenopus egg extracts from phosphoinositide-containing lipid bilayers. Methods in Cell Biology 128, 125–147.

Wang, H, Lo, W-T, Žagar, AV, Gulluni, F, Lehmann, M, Scapozza, L, Haucke, V, and Vadas, O (2018). Autoregulation of Class II Alpha PI3K Activity by Its Lipid-Binding PX-C2 Domain Module. Mol Cell 71, 343–351.e4.

Wu, M, Huang, B, Graham, M, Raimondi, A, Heuser, JE, Zhuang, X, and Camilli, PD (2010). Coupling between clathrin-dependent endocytic budding and F-BAR-dependent tubulation in a cell-free system. Nat Cell Biol 12, 902–908.

Wühr, M, Freeman, RM, Presler, M, Horb, ME, Peshkin, L, Gygi, SP, and Kirschner, MW (2014). Deep Proteomics of the Xenopus laevis Egg using an mRNA-Derived Reference Database. Curr Biol 24, 1467–1475.

Yang, S, Huang, F-K, Huang, J, Chen, S, Jakoncic, J, Leo-Macias, A, Diaz-Avalos, R, Chen, L, Zhang, JJ, and Huang, X-Y (2013). Molecular Mechanism of Fascin Function in Filopodial Formation*. J Biol Chem 288, 274–284.

Yarar, D, Surka, MC, Leonard, MC, and Schmid, SL (2008). SNX9 Activities are Regulated by Multiple Phosphoinositides Through both PX and BAR Domains. Traffic 9, 133–146.

Yarar, D, Waterman-Storer, CM, Schmid, SL, and Biology, TSRID of C (2007). SNX9 Couples Actin Assembly to Phosphoinositide Signals and Is Required for Membrane Remodeling during Endocytosis. Developmental Cell 13, 43–56.

Zecchini, V, Paupe, V, Herranz-Montoya, I, Janssen, J, Wortel, IMN, Morris, JL, Ferguson, A, Chowdury, SR, Segarra-Mondejar, M, Costa, ASH, et al. (2023). Fumarate induces vesicular release of mtDNA to drive innate immunity. Nature 615, 499–506.

Zhang, R, Lee, DM, Jimah, JR, Gerassimov, N, Yang, C, Kim, S, Luvsanjav, D, Winkelman, J, Mettlen, M, Abrams, ME, et al. (2020). Dynamin regulates the dynamics and mechanical strength of the actin cytoskeleton as a multifilament actin-bundling protein. Nat Cell Biol 22, 674–688.

Zheng, S, Wolff, G, Greenan, G, Chen, Z, Faas, FGA, Bárcena, M, Koster, AJ, Cheng, Y, and Agard, DA (2022). AreTomo: An integrated software package for automated marker-free, motion-corrected cryo-electron tomographic alignment and reconstruction. J Struct Biol: X 6, 100068.

